# Synthetic fibrous hydrogels as a platform to decipher cell-matrix mechanical interactions

**DOI:** 10.1101/2022.08.24.505064

**Authors:** Hongbo Yuan, Kaizheng Liu, Mar Cóndor, Jorge Barrasa-Fano, Boris Louis, Johannes Vandaele, Paula de Almeida, Quinten Coucke, Wen Chen, Egbert Oosterwijk, Chenfen Xing, Hans Van Oosterwyck, Paul H. J. Kouwer, Susana Rocha

**Affiliations:** Key Laboratory of Molecular Biophysics of Hebei Province, Institute of Biophysics, School of Health Sciences and Biomedical Engineering, Hebei University of Technology, Tianjin 300401, P. R. China; Molecular Imaging and Photonics, Chemistry Department, KU Leuven, Celestijnenlaan 200F, 3001 Heverlee, Belgium; Institute for Molecules and Materials, Radboud University, 6525 AJ Nijmegen, The Netherlands; Institute of Biomedicine and Biotechnology, Shenzhen Institutes of Advanced Technology, Chinese Academy of Sciences, Shenzhen, 518055, P. R. China; Department of Mechanical Engineering, Biomechanics section, KU Leuven, Belgium; Division of Chemical Physics and NanoLund, Lund University, Sweden; Radboud Institute for Molecular Life Sciences, Department of Urology, Radboud University Medical Center, 6500 HB Nijmegen, The Netherlands; Prometheus Division of Skeletal Tissue Engineering, KU Leuven, Belgium

## Abstract

The interactions between cells and their direct environment are crucial for cell fate but biochemically and mechanically highly complex, and therefore, poorly understood. Despite recent advances that exposed the impact of a range of different factors, real progress remains challenging, since appropriate controllable matrices and quantitative analysis techniques that cover a range of time and length scales are unavailable. Here, we use a synthetic fibrous hydrogel with nonlinear mechanics to mimic and tailor the bi-directional cell-matrix interactions. Using advanced microscopy-based approaches, we acquire a comprehensive picture of how cellular traction forces, fiber remodeling, matrix stiffening, matrix properties and cellular behavior interact, highlighting for instance, the importance of a fibrous architecture and nonlinear mechanics of the matrix. Complete mapping of cell-matrix interactions at the cellular length scale provides indispensable information for the rational design of biomimetic materials to recreate realistic in vitro cell environments.

---

Cells are intimately linked to their surrounding extracellular matrix (ECM); they receive chemical cues and constantly probe and respond to external mechanical properties. Simultaneously, cells can actively modify the mechanical properties of their environment by secreting or remodeling ECM components, which gives rise to a complex mechanical reciprocity that plays a key role in cell migration and development, tissue regeneration and homeostasis, and pathologic changes^1,2^. Cellular actomyosin related contraction generates forces that propagate over distances many times the cell diameter, allowing cell-cell mechanical communication and rapid transmission of signals at a long distance^3–5^. Additionally, cellular contraction leads to large stiffness gradients in the 3D matrix, i.e. the ECM closer to the cells becomes substantially stiffer, due to matrix densification and a stress-induced stiffening response of the matrix^6–13^. As the latter is commonly only found in biological matrices, cell-induced matrix stiffening and force propagation over large distances have only been observed in collagen and fibrin matrices^14,15^. Although biocompatible, these matrices are relatively difficult to tailor^16^, particularly without modifying the protein concentration, which inherently changes other matrix characteristics too (e.g. pore size, concentration of adhesion ligands). Moreover, such biological matrices are sensitive to enzymatic degradation, which will change matrix properties in time.

Synthetic hydrogels are uniquely suited to decouple biochemical and mechanical cell-matrix interactions. In this work, we recreated the mechanical cell-matrix interactions observed in biological materials in a fully synthetic hydrogel based on polyisocyanides (PICs), which allowed us to study the reciprocal cell-matrix interactions in a very selective and controlled fashion. Using a combination of 3D displacement microscopy, equivalent to traction force microscopy (TFM), and live cell rheology, we delineated characteristic time and length scales of force propagation and matrix stiffening, key information for the understanding cell-matrix mechanical interactions and rational design of the next generation of biomimetic materials.

## Cell-mediated mechanical remodeling

The PIC-based hydrogels used here combine the structural and mechanical characteristics of a biological gel, with the tailorability of a synthetic gel^17–20^. Decoration with the generic cell-adhesive peptide GRGDS (PIC^*RGD*^) makes the material particularly suitable as a matrix for the 3D culture of different cells and organoids^21–23^, where control over (mechanical) matrix properties is used to direct cell fate, such as stem cell differentiation^24^, cell morphogenesis^21,25^ and the secretome^26^. Live cell imaging of human adipose-derived stem cells (hASCs) in fluorescently labelled PIC^*RGD*+^ (see Methods section and Supplementary Fig. S1 for details) shows cell migration and matrix remodeling, as indicated by the increased fluorescence intensity of the matrix close to the cell (Fig. 1a,b, Supplementary Movie 1). In addition, it is possible to observe channels formed by migrating cells (Fig. 1a, dotted lines). Since the PIC network is not chemically crosslinked but held together with noncovalent interactions^17^, we hypothesize that the observed remodeling is purely mechanical and results from mechanical cell-matrix interactions, in line with the migration mechanism described for alginate-based gels^9^. Plasticity of the matrix ensures that the formed channels remain in the material. Further zooming in on the early (24 h after incubation) remodeling process taking place around the cell, the images acquired show a radially oriented fiber alignment (Fig. 1b, white arrows), fiber densification and clear stress propagation tracks throughout the matrix.

**Fig. 1.**
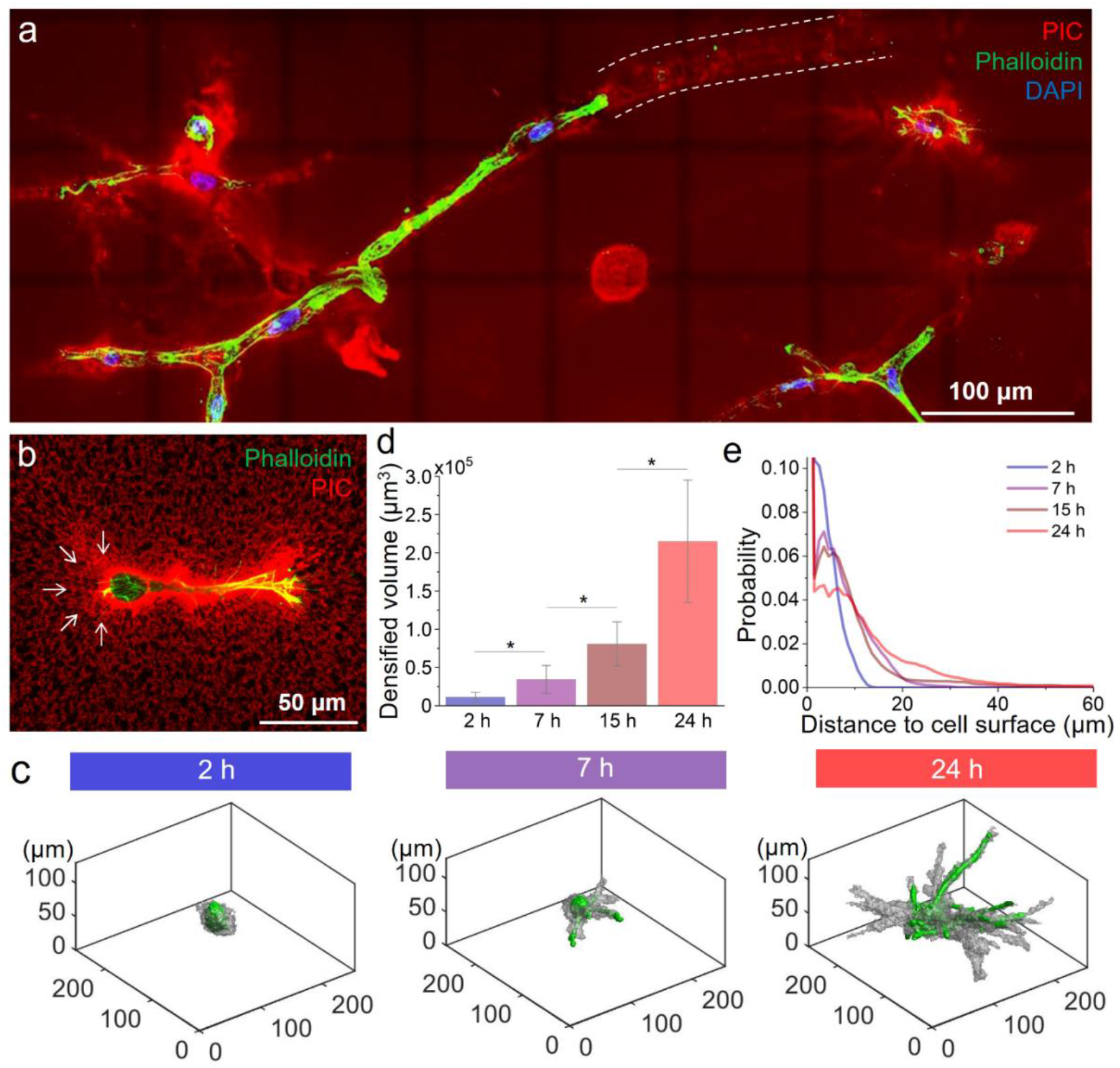
Cell-mediated mechanical remodeling in PIC matrices. (a) Maximum projection image of hASCs in a fluorescently labelled 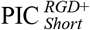 gel 7 days after encapsulation (different PIC gels are discussed in a next section). White dotted lines indicate the tunnels formed by cell migration that remain in the PIC matrix. (b) Higher resolution fluorescence image of a hASC in 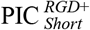 network 24 h after incubation. The white arrows indicate areas of fiber recruitment of PIC with radial orientation. Red: PIC fibers. Green: phalloidin. Blue: DAPI. (c) 3D render images of representative cells, 2, 7 and 24 h after encapsulation, displaying the cell surface (green) and the volume of remodeled matrix (grey). The remodeled matrix was determined using an intensity-based threshold (details in the methods section). (d) Evolution of the densified fiber volume through time (N = 5, * indicates p<0.05, by One-way ANOVA Tukey’s test). (e) Distribution of the fiber remodeling range with incubation time. The PIC matrix used in these experiments were 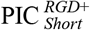 with short polymer lengths at 1.0 mg mL^−1^, yielding gels with a low critical stress and high sensitivity to cellular forces (parameters in Table 1).

To quantitatively investigate the dynamics and extent of cell-induced fiber remodeling with time, regions displaying fiber densification were defined using an intensity-based threshold, following the approach described by Lesman and coworkers^14^ (Fig. 1c,d). Two hours after cell encapsulation, the extent of remodeling is almost negligible but after that, the matrix volume that shows cell-induced fiber densification gradually increases and the distance of remodeled matrix, as measured from the cell surface, increases from less than 10 µm at 2 h to more than 50 µm after 24 h (Fig. 1e, Supplementary Fig. S2). Despite being a common phenomenon in natural matrices, this is the first time that such long-range mechanical remodeling of matrix fibers is reported in a fully synthetic material.

## Cell-induced fiber displacements and force propagation

To evaluate cellular force-induced matrix displacements and remodeling, we monitor changes in the fiber architecture of the PIC gel after the removal of the contractile forces by addition of the actin disrupting agent cytochalasin D (cytoD)^27,28^. In contrast to conventional TFM, there is no need to add particles to indirectly monitor matrix displacements: the fluorescently labelled PIC gels allow direct bead-free tracking (Supplementary Fig. S1).

Three-dimensional cultures of hASCs containing a fluorescent dye (CellTracker™ Green) encapsulated in the GRGDS-decorated and fluorescently labelled PIC hydrogel were imaged after 2, 7, 15 or 24 h of incubation. The cells were then relaxed by adding cytoD and imaged again (Supplementary Fig. S3 and Movie 2). Fiber displacements between the stressed and relaxed state of the cell were measured using the open-source 3D TFM toolbox TFMLAB, to generate matrix displacement maps. The arrows in the displacement maps indicate the direction and magnitude of the measured displacements (Fig. 2a-d, details in the Methods Section)^29^. The arrows in the displacement maps indicate the direction and amplitude of the detected displacements.

**Fig. 2.**
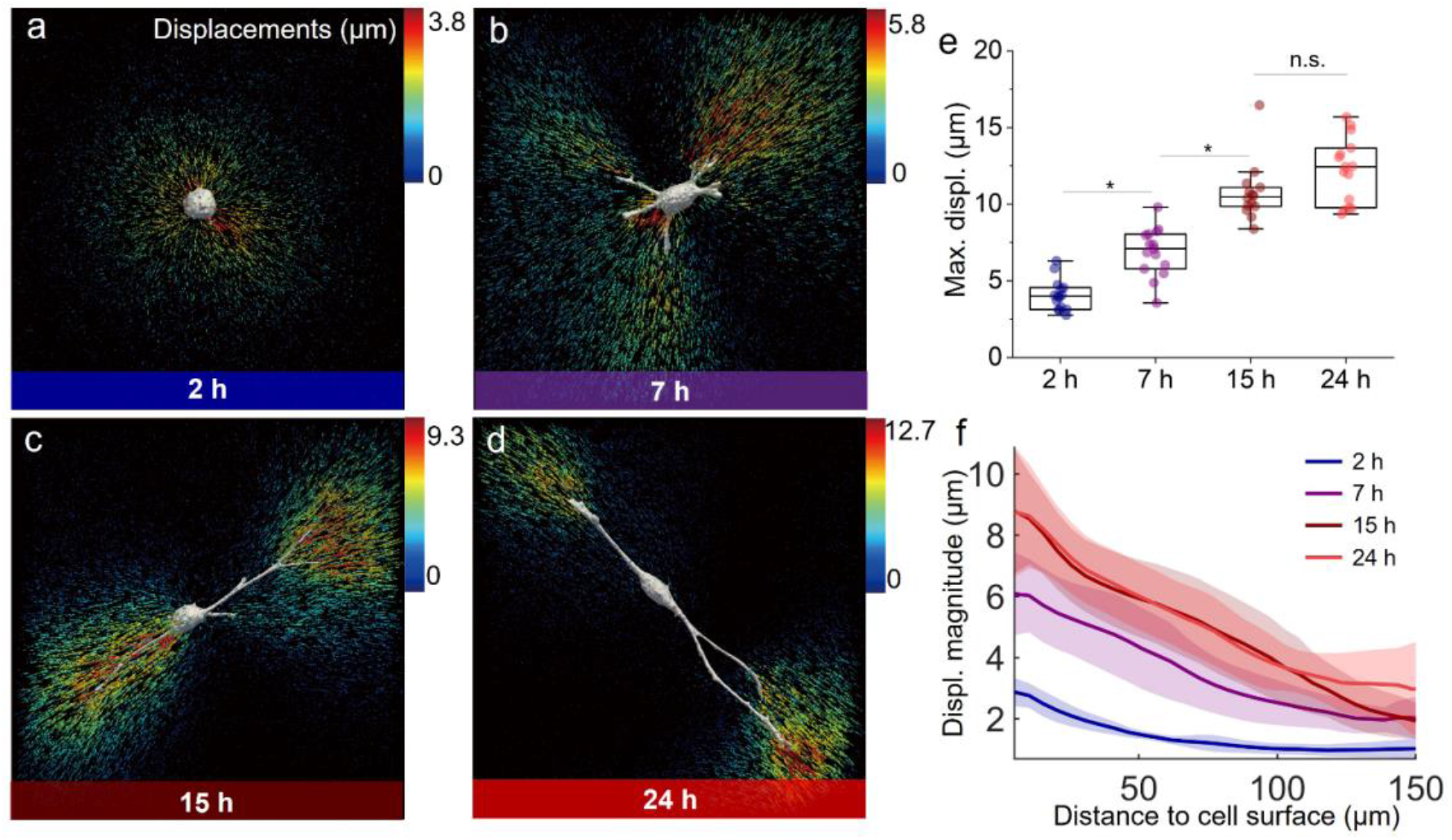
Cellular force-generated fiber displacements and transmission range in PIC matrices. (a-d) 3D displacement field induced by hASCs encapsulated in 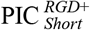 gel after the addition of cytoD at different time points (2, 7, 15 and 24 h respectively). The arrows indicate the calculated displacement of the PIC fiber network and are color-coded according to the magnitude of the local displacement vector (in μm). Notice the different scales for a-d. (e) The maximum displacement (Max. displ.) plotted for each cell at different incubation times (N = 12-15, * indicates p<0.05, by One-way ANOVA Tukey’s test). (f) Plots of the amplitude of the fiber displacements as a function of distance to cell surface, at different time points. 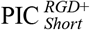 with a default concentration 1.0 mg mL^−1^ was used (Table 1).

The measurement of cell-induced matrix displacements with time provides quantitative information on the dynamics of cell-matrix interactions^31^. Two hours after encapsulation, the cells are mostly spherical and apply uniform, radially oriented traction forces on its surroundings, pulling the fiber network towards the center of the cell (Fig. 2a), which is in line with the observed patterns of fiber remodeling. With time, cells spread and develop protrusions (Fig. 2b-d), which gradually become longer and slenderer. We find that the largest matrix deformations are present at the tips of these protrusions. To quantify the evolution of matrix displacements through time, the maximum fiber displacement for each cell was determined (Fig. 2e). During the first 15 hours in culture, the average maximum displacement gradually increases over time, until reaching a plateau where cells are considered stable in the gel as no significant differences in displacements can be detected after 15 h of incubation.

The range of cell-induced displacements into the fibrous matrix was calculated by plotting fiber displacements as a function of the distance from the cell surface (Fig. 2f). After 7 h in culture, we detect fiber displacements as far as 150 μm from the cell surface^32^, which is in strong contrast to values that have been reported using linear elastic materials (such as polyethylene glycol or polyacrylamide gels) where the displacement amplitudes rapidly decay within a distance at the same order of the cell diameter (approximately 20 μm)^33,34^.

In addition, the formation of bundles of aligned fibers between cells in close proximity was also observed (Supplementary Fig. S4), a phenomenon that was reported only in biological matrices like reconstituted collagen^35^ and fibrin^14^ gels. Such aligned fiber structures are known to be involved in long-range mechano-communication and molecular transport between cells in 3D environments^4^ and required the unique, nonlinear responses of fibrous networks present in the natural ECM^3^.

## Cell-mediated matrix stiffening and plastic remodeling

Cell-induced matrix stiffening, as earlier observed in natural biopolymer networks such as collagen, fibrin and Matrigel^6,7^ is usually attributed to the combination of contractile cellular forces and the nonlinear elastic behavior of the materials. The contribution of matrix remodeling, including fiber densification is often overlooked. Here, we combine live-cell rheological tests and fluorescence image analysis to unravel the role of cellular forces and fiber remodeling in cell-induced matrix stiffening.

The storage modulus *G*′ of PIC gels containing hASCs appears as a rising curve that plateaus after approximately 15 h (Fig. 3a, red dots). The final storage modulus at 24 h is about three times higher than that of the gel without cells (grey squares) and remains relatively constant up to 48 hours (Supplementary Fig. S5a). Lower cell densities reduce the stiffening effect (Supplementary Fig. S5b). PEG microspheres (20 μm diameter) or even less-contractile HeLa cells do not induce stiffening but rather a small decrease in *G*′ (Fig. 3b), as a result of reduced entanglements in the matrix^36^. Analogously, when hASCs are encapsulated in GRGDS-free PIC gels (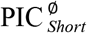 in Table 1), network softening is observed (Fig. 3b), which confirms that the observed increase requires contractile cells, cell-matrix interactions and a strain stiffening matrix. The time scale of matrix deformation observed using fluorescence microscopy (Fig. 2e) coincides with the macroscopic stiffening response measured by rheology, suggesting that both effects are caused by cellular tractions.

**Table 1.** Characterization of the library of PIC gels, collagen and Matrigel matrices used.

| Matrix | $L_c$<br>(nm) <sup>[a]</sup> | $c$<br>(mg mL<sup>-1</sup>) | GRGDS<br>density (μM) | Shear<br>modulus $G'$<br>(Pa) <sup>[b]</sup> | Critical stress<br>$\sigma_c$ (Pa) <sup>[b]</sup> |
| --- | --- | --- | --- | --- | --- |
| PIC<sup>∅</sup><sub>Short</sub> | 154 | 1.0 | — | 47 | 10.6 |
| PIC<sup>RGD</sup><sub>Short</sub> | 154 | 1.0 | 3.7 | 46 | 8.6 |
| PIC<sup>RGD+</sup><sub>Short</sub> | 154 | 1.0 | 31.4 | 45 | 7.7 |
| PIC<sup>RGD+</sup><sub>Long</sub> | 229 | 1.0 | 31.4 | 93 | 19.9 |
| 1.5×PIC<sup>RGD+</sup><sub>Short</sub> | 154 | 1.5 | 47.1 | 89 | 11.6 |
| Collagen | n/a <sup>[c]</sup> | 1.2 | n.d. <sup>[d]</sup> | 60 | 3.5 |
| Matrigel | n/a <sup>[c]</sup> | 8-11 <sup>[e]</sup> | n.d. <sup>[d]</sup> | 28 | 6.9 |

Note: <sup>[a]</sup> The polymer contour length, $L_c$ , was calculated from the viscosity-based molecular weight; <sup>[b]</sup> mechanical properties were measured in $\alpha$ MEM at 37 °C, in the absence of cells; <sup>[c]</sup> not applicable; <sup>[d]</sup> not determined; <sup>[e]</sup> Typical protein concentrations for Corning Matrigel matrix are between 8 to 11 mg mL<sup>-1</sup>.

**Fig. 3.**
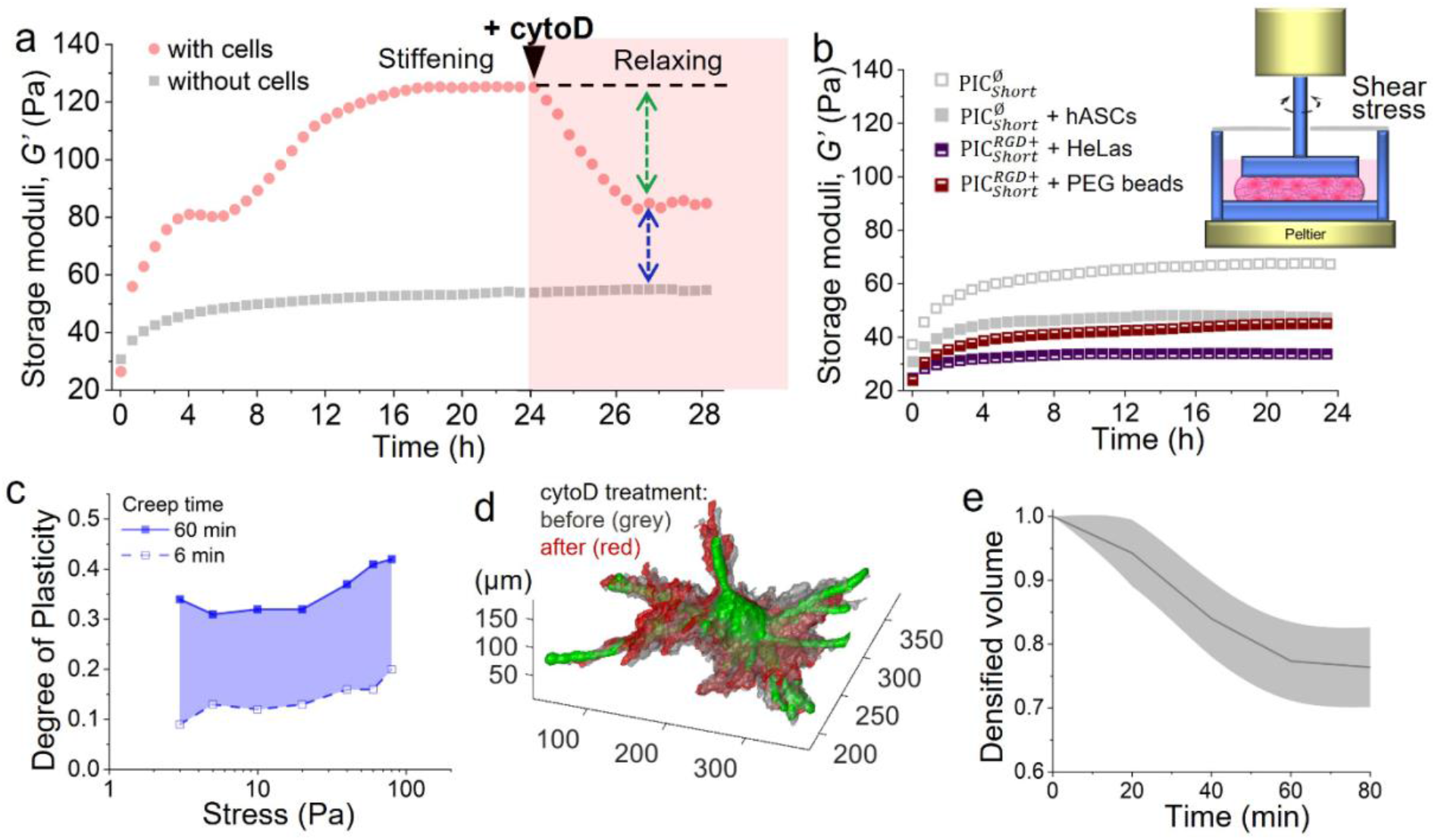
Cell-mediated matrix stiffening and plastic remodeling in PIC matrices. (a) The storage moduli *G*′ of 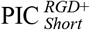 gels with (red dots) or without hASCs (grey squares) were monitored in real time through live-cell rheological measurement. Left panel indicates the cell-induced stiffening behavior for the first 24 h before adding cytoD, and the right panel in pink indicates the relaxing effect for an additional 4 h after adding cytoD (5 μM). The green arrow indicates the decreased in *G*’ caused by the addition of cytoD while the blue arrow indicates the irreversible change in *G*’ (compared to the sample without cells). (b) Stiffness of cell adhesion peptide-free gels 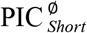 and 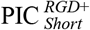 with cells and PEG microspheres as a function of time. hASCs (contractile) and HeLa cells (less-contractile) were used as model cells. Inset: Illustration of the setup used for live-cell rheology. A cold solution of PIC with or without cells was placed inside a shear rheology setup with a petri dish-like plate geometry. The solution was warmed to 37 °C to form a gel, and then warm CO_2_-independent medium was added into the dish. HeLas and hASCs in RGD-free gels maintain a spherical morphology (Supplementary Fig. S6). (c) Degree of plasticity of 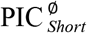 gels as a function of stress with different creep time. Solid line: 60 min, dashed line: 6 min. At increasing timescales, PIC gels exhibit higher degrees of plasticity. (d) Representative 3D rendering of the remodeled matrix before (grey) and 1 h after (red) adding cytoD. The initial image was acquired 24 h after incubation. The cell surface is depicted in green. (e) Evolution of the densified fiber volume over time after adding cytoD (N = 5). Both 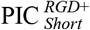 and 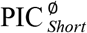 polymers were used at a concentration of 1.0 mg mL^−1^.

Cell contraction, and the resulting macroscopic stiffening response, were terminated by the addition of cytoD, which decreased *G*′ but not to the value without cells (Fig.3a, red dots). We hypothesize that the remaining stiffening is caused by plasticity-mediated matrix remodeling, i.e. the architectural reorganization of the polymer network induced by cellular remodeling. Due to the plasticity of the PIC network, full recovery of the remodeled structure after cell relaxation is prevented. Plasticity is qualitatively inferred from the cell migration channels in the fluorescence images (Fig. 1a). We then studied if plastic remodeling of the matrix directly affects the mechanical properties. We performed a series of macroscopic creep and recovery tests (Supplementary Fig. S7) and found that while the degree of plasticity increases with applied stress (Fig. 3c), the stiffness of the matrix remains constant for all plastic deformations.

To quantify the plastic character of the cell-induced matrix remodeling at the micrometer scale, we traced and analyzed time-lapse fluorescence images of the fiber network after inhibiting cellular contraction. CytoD addition reduced the length of the cellular protrusions as the cell reached the stress-free state (Fig. 3d,e). Simultaneously, the volume of densified PIC matrix gradually decreased, until approximately 25% of the initial densified volume, which indicates that a large fraction of the matrix surrounding the cell is deformed plastically and irreversibly (~75%, see Fig. 3e). We attribute the residual macroscopically observed stiffening after cytoD addition to the (local) permanently remodeled architecture of the PIC matrix. Overall, we conclude that the observed matrix stiffening has two origins, both induced by cellular contraction: a fast nonlinear strain-stiffening response that is fully reversible and a slower and irreversible plastic remodeling response that includes fiber densification and alignment.

## Nonlinear mechanics of the fibrous network dominates the cell-matrix mechanical interactions

The nonlinear mechanics of PIC gels are readily tuned by the polymer concentration, *c*, and also at constant *c* by the polymer contour length^19^ (*L*_c_). For instance, for gels of shorter polymers the critical stress *σ*_c_ (the onset of nonlinear stiffening) decreases, which gives rise to a material that is more sensitive to stress^19^. Fluorescence microscopy on labelled PIC gels, however showed that the fiber architecture at the micrometer scale depends on *c*, but is not affected by *L*_*c*_ ^37^, which allows one to tune the mechanical properties without affecting the gel microstructure.

Here we exploited the tailorability of the PIC gel to learn more about cell-matrix interactions. hASCs were encapsulated (3D) in three different PIC gels. Two types of gel were comprised of short PIC polymers, with medium and high GRGDS densities (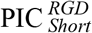 and 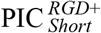, respectively; and a third gel, made of long PIC polymers, with a high GRGDS density 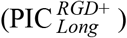. Gels prepared from short polymer exhibit a low critical stress, which indicates a higher sensitivity to mechanical or cell-induced stress. In contrast, long PIC polymers form gels with a higher critical stress, presenting lower mechanical sensitivity. The properties of the gels used are summarized in Table 1.

Twenty-four hours after encapsulation, live cell imaging was performed to evaluate cell morphology, fiber displacements and the degree of matrix remodeling (Fig. 4). In line with previous work^21^, we found that the spreading of hASCs is stronger in RGD-rich soft matrices 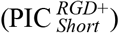, characterized by the long and numerous slender protrusions and an elongated morphology (Fig. 4a and Supplementary Fig. S8). Spreading reduces with decreased concentration of adhesion ligands or in a stiffer matrix (Fig. 4b,c). Image analysis of 12-15 representative cells confirms that cells in 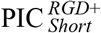 form more protrusions (Fig. 4d) and increased levels of fiber displacement and propagation distance (Fig. 4e,f).

**Fig. 4.**
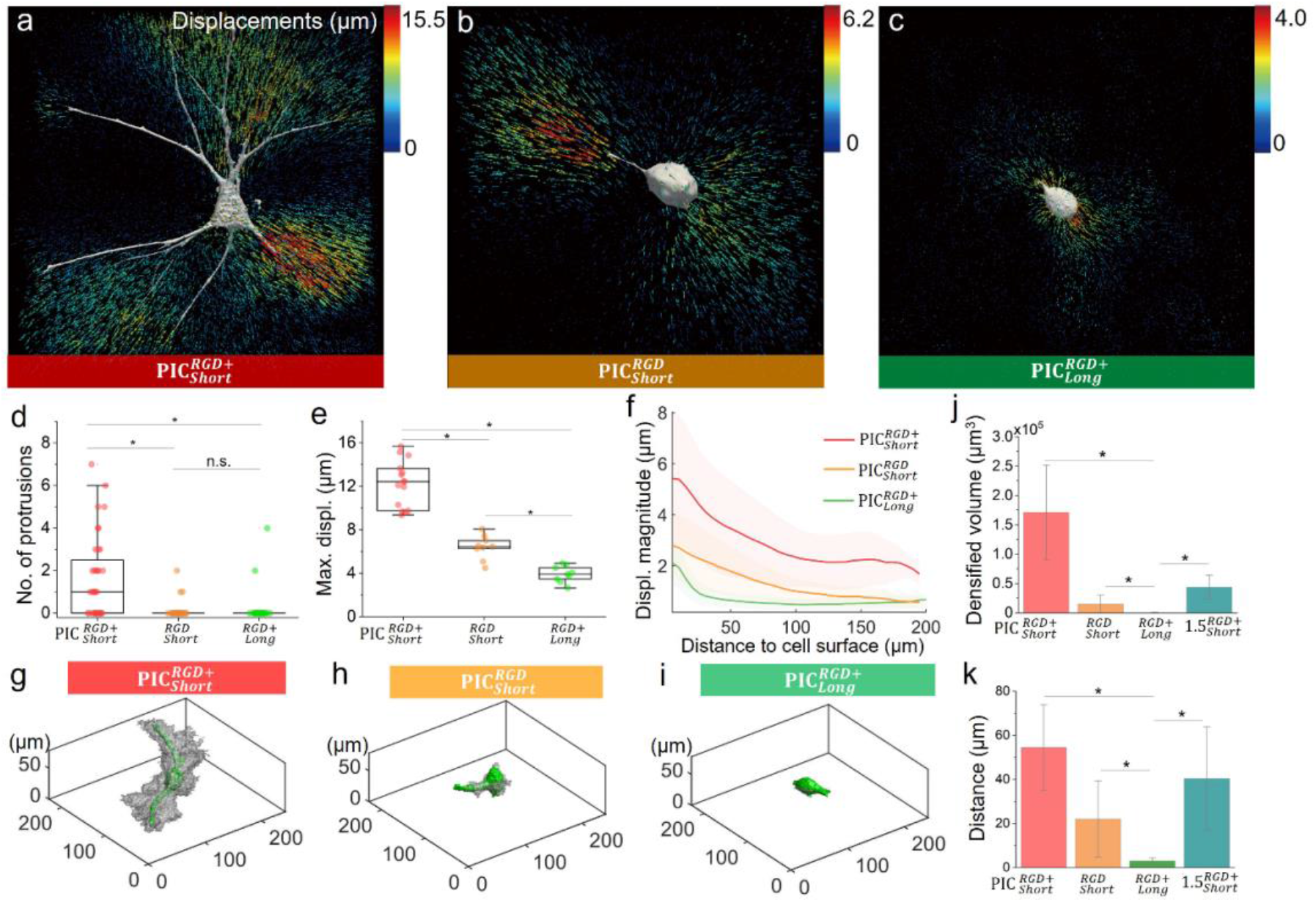
Influence of the bio/mechanical properties on cell-matrix mechanical interactions. hASCs-induced 3D matrix displacements in 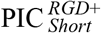 (a), 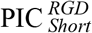 (b), and 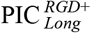 (c) gels at 24 hours after incubation. Arrows indicate displacement directions and are color coded for magnitudes (notice the different scales for a–c). (d) Number of protrusions in hASCs in different PIC gels (N = 20-25). (e) Maximum cell-induced matrix displacement (Max. displ.) in the different PIC gels (N = 12-15). (f) Plots of fiber displacements as a function of distance from the cell surface (N = 8-10). (g-i) Plots of a representative cell in different PIC gels, displaying the cell body (green) and the volume of remodeled matrix (grey). The densified volumes were detected using an intensity based adaptive threshold (details in the Methods section). (j) Densified fiber volume in different PIC matrices (N = 5). (k) Maximum distance of the detected edge of fiber densification for the different PIC matrices (N = 5). 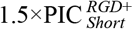 refers to the sample prepared with short 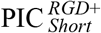 polymers but with higher concentration (1.5 mg mL^−1^) compared to the default condition in other experiments (1.0 mg mL^−1^), see the detailed parameters in Table 1. * indicates p<0.05, by One-way ANOVA Tukey’s test.

Matrix stiffening as a result of plastic remodeling is directly related to the volume of the remodeled matrix around the cells. Further image analysis allows us to generate a full 3D displacement field around the hASCs in each matrix (Fig. 4g-i, the averaged data of densified volume and distance to the cell surface is shown in panels j and k). The matrix displays more pronounced remodeling in 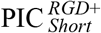 and much less for the gels containing less RGD or stiffer gels with the same polymer concentration but longer chains. In addition, we note that while the amplitude of the matrix deformation in 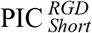 was lower compared to 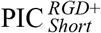, the range of propagation and decay of the cellular force-induced displacements was similar in both gels (Fig. 4e,f). In contrast, in the 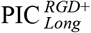 gels, fiber displacements were only detected in the pericellular region, rapidly decreasing within 30 μm from the cell surface, which indicates that force transmission range mainly depends on the mechanical properties of the matrix and is less affected by the number of cell-matrix adhesion sites.

The differences observed between hydrogels composed of long and short polymer chains can be linked to higher critical stress or higher storage modulus (Table 1). To evaluate the contribution of the nonlinear mechanics, we prepared a gel of 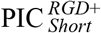 at a slightly higher polymer concentration (*c* = 1.5 mg mL^−1^, labelled as 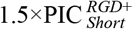 in Table 1), with a shear modulus *G*′ comparable to 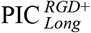 (at *c* = 1.0 mg mL^−1^) but a lower critical stress (11.6 versus 19.9 Pa, see Table 1). Fluorescence microscopy after 24 h incubation (Supplementary Fig. S9) shows that hASCs embedded in 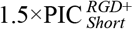 aligned and densified the matrix. While the densified volume was smaller than in 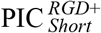, it was still significantly larger than that in the 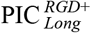 gels (Fig. 4j,k), which suggests that the nonlinear mechanics of PIC gels plays an important role in the cell-matrix interactions. We propose that for the long chain PIC gels, the threshold stress at which the polymers slide and merge to remodel is beyond the range of hASCs-generated forces and therefore the amount of fiber remodeling is greatly reduced for increasing values of critical stress.

## Comparison with biological matrices

To analyze the biological relevance of our analysis on synthetic matrices, we benchmark our results to cell-induced matrix displacements in natural matrices. hASCs were encapsulated in collagen and Matrigel matrices; we aimed to roughly match the linear and nonlinear mechanical properties of the biopolymer networks to 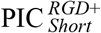, despite the strong differences in strain stiffening effects in the three matrices (Table 1).

Twenty-four hours after encapsulation, hASCs displayed a well-spread morphology in both PIC and collagen matrices, while spreading was less in Matrigel (Fig. 5a-c). Cells in PIC and collagen show a similar cellular sphericity, volume, number of protrusions and protrusion length, while in Matrigel the cells are less spread, showing smaller protrusions (Fig. 5d-g). Displacement analysis reveals that the fiber displacement field observed in PIC hydrogels is close to that observed in collagen (Fig. 5a-b) with no significant differences on the calculated maximum displacement (Fig. 5h). Remarkably, the force propagation was also similar in both matrices (Fig. 5i); both PIC and collagen gels support a long-range fiber displacement propagation, reaching over hundred microns from the cell surface. The fewer and shorter protrusions in Matrigel generated a much smaller and more local displacements (note the different color scales in Fig. 5a-c).

**Fig. 5.**
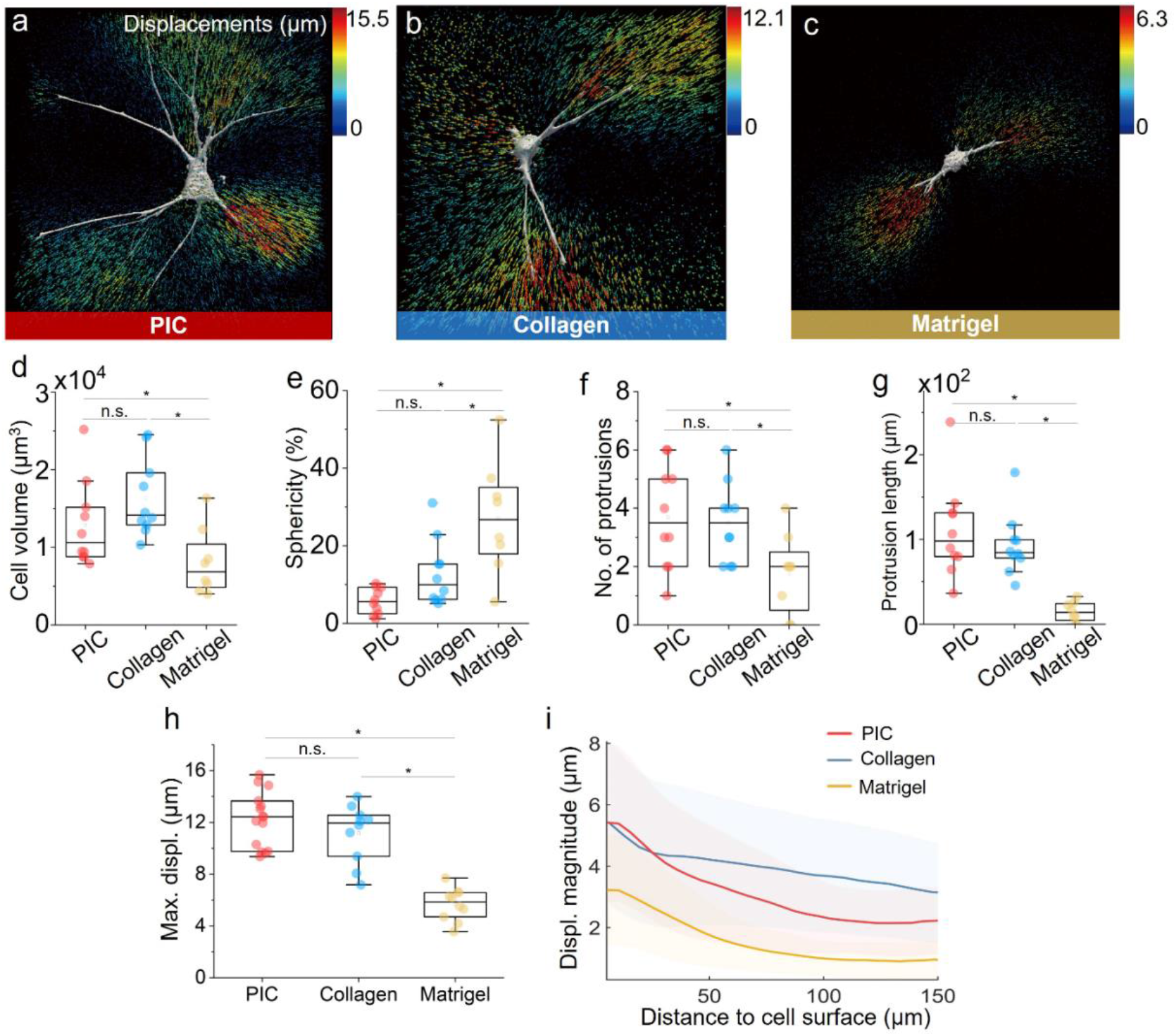
Comparison of cell behavior and cell-matrix mechanical interactions between PIC and biological matrices. hASCs induced 3D matrix displacement fields in PIC (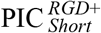, 1.0 mg mL^−1^) gel (a), collagen type I (1.2 mg mL^−1^, b) and Matrigel (70%, c). The colour of the arrows indicates the magnitude of the local displacement vector. Comparison of cell morphology in different matrices after 24 hours incubation, including cell volume (d) and sphericity (e), protrusion number (f) and length (g). (h) Maximum cell-induced matrix displacement (Max. displ.) in different matrices. (i) Plots of matrix displacements as a function of distance to cell surface. N = 8-12, * indicates p<0.05, by One-way ANOVA Tukey’s test.

In general, cell spreading is affected by matrix mechanics, network porosity and integrin-mediated cell-matrix interactions^38^. Similar to collagen, PIC gels present a fibrous architecture with pore size in the micrometer range^37^. The α_5_β_1_ integrin dimer, highly abundant in hASCs, binds well to RGD and to collagen type I^39,40^, which gives rise to pronounced cellular spreading and an extensive matrix remodeling. Matrigel, on the other hand, exhibits a much denser and sterically constrained matrix^41^, with laminin as the main component, which interacts primarily with α_3_β_1_ integrins (that are less expressed in hASCs)^39,40^. We observe that the propagation of the cellular forces through the Matrigel matrix resembles the decay observed in RGD-reduced PIC gels (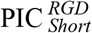, Fig. 4e and 5i), which seems to suggest that the reduced cell spreading and smaller fiber displacements observed in Matrigel is a consequence of lesser cell-matrix interactions.

## Discussion

Over the last two decades, a plethora of studies have demonstrated unambiguously that the mechanical properties of the cell’s environment determine its fate. While it is often easy to measure these properties macroscopically, it remains very difficult to analyze them in 3D samples at the micron scale, i.e., at the cellular length scale, particularly in fibrous matrices such as collagen, which are not homogeneous at this scale. In this manuscript, we showed that cells bind to a matrix and apply contractile forces to it, actively modifying their environment, even when it is nondegradable. With a combination of macroscopic (live cell) mechanical analysis and microscopic 3D displacement measurements, we were able to map how cellular contractile forces lead to remodeling and stiffening through two distinct mechanisms. At short timescales, cellular contraction deforms the matrix, which instantaneously results in a stiffening response due to the nonlinear strain stiffening properties of the matrix, which is directly linked to its fibrous architecture. This effect is temporary, i.e., as the stress is removed, for instance by cytoD addition, or at longer timescales by viscoelastic dissipation^16^, matrix stiffening disappears.

At longer timescales, cells are able to remodel their matrix, densifying and aligning it. In fibrous matrices, the results of remodeling are not only noticed close to the cell, but the effect can reach out hundreds of microns into the matrix, which sets up a mechanism for easy long-range cellular communication. The high plasticity of the matrix makes remodeling (partially) permanent and even after removal of cellular tractions, the local matrix densification and alignment remain, leading to matrix changes that can be macroscopically traced.

To study biologically relevant cell-matrix interactions, one ideally incorporates as many as possible characteristics of the *in vivo* microenvironment. From our work, it becomes increasingly clear that key design features include a fibrous architecture (preferably with associated nonlinear mechanical properties) and sufficient cell-matrix interactions to achieve a reliable traction response. Most likely, the latter requires optimization for each cell type^43^. In that respect, synthetic materials are particularly attractive as matrices because of their high customizability, both mechanically and biochemically, which can also be used to drive desired effects. When studying at longer timescales, also effects of matrix degradation and the deposition may be considered.

Taken together, the interdisciplinary combination of a customizable biomimetic material and advanced fluorescent microscopy techniques has the potential to become a highly valuable cell biology toolbox to study reciprocal cell-matrix interaction in dilute systems as well as more complex heterocellular systems.

## Online methods

### Polymer synthesis and functionalization

#### PIC polymer synthesis

Polyisocyanides were synthesized using an established protocol as reported previously^21^. The polymer contour length, *L*_C_, was controlled by adjusting the total monomer to catalyst ratio and estimated from the viscosity averaged molecular weight. All azide-modified polymers carried 3.3% azide monomers and 96.7% methoxy-terminated monomers.

#### Peptide conjugation

A linear GRGDS peptide with BCN (bicyclo[6.1.0]nonyne) was used. The linker between the peptide and the BCN reactive handle was PPPSG[Abz]SG, where Abz is 4-aminobenzoic acid. Peptides were purchased from Pepsan, the Netherlands. The azide-decorated polymer was first dissolved in acetonitrile (2.5 mg/mL), and the appropriate volume of the peptide solution was added (6.0 mg/mL in DMSO). The solution was stirred for 24 h at room temperature. Next, the polymer–peptide conjugates were precipitated in diisopropyl ether, collected by centrifugation, and air-dried for 2-3 days.

#### Fluorescence labelling

To fluorescently label the polymers, a red-emitting fluorescence dye Atto647N with covalently grafted. Atto647N-NHS ester was conjugated with DBCO-PEG4-amine (Sigma Aldrich), yielding DBCO-Atto647N. This compound was added to the polymer solution in a ratio of 1 mg polymer: 0.5 nmol of dye and incubated for 5 minutes on ice, and then used for cell encapsulation.

### Cell culture

#### Cell Passage

Human adipose stem cells (hASCs) were isolated from patients with informed consent and grown in α-MEM medium (Gibco, Thermo Fisher, USA) supplemented with 10% fetal bovine serum (FCS) (Gibco, Thermo Fisher, USA) and 1% Penicillin/Streptomycin (10,000 Units/mL Pen, 10,000 μg/mL Strep, Gibco, Thermo Fisher). The cells from the same patient sample and the same passage number (p = 6) were used for experiments. The HeLa cells were cultured in RMPI 1640 media (Gibco, Thermo Fisher) supplemented with 10% fetal bovine serum (FCS) (Gibco, Thermo Fisher, USA) and 1% *L*-Glutamine (Gibco, ThermoFisher, USA). Cells were incubated at 37 °C, 5% CO_2_.

#### Encapsulation in PIC gels

Cells were encapsulated in PIC gels by mixing a cell suspension and polymer solution (v/v = 1:1) at lower temperature (on ice) and warming up to 37 °C in an incubator. The initial PIC polymer solution was 2.0 mg/mL, resulting in a final concentration of 1.0 mg/mL (expect when specified otherwise). The final cell density was 10^6^ cells/mL for live cell rheology experiments, 8 × 10^4^ cells/mL for the analysis of cell morphology, and 4 × 10^4^ cells/mL for TFM. Imaging was performed using angiogenesis plates (Ibidi).

#### Encapsulation in collagen gels

The collagen gels used in this work were prepared using a protocol previously published^44^. Briefly, to prepare 310 μl of 2.4 mg/ml collagen type I hydrogels, we mixed 125 μl of rat tail collagen (Collagen R, 2 mg/ml, Matrix Bioscience), 125 μl Bovine skin collagen (Collagen G, 4 mg/ml, Matrix Bioscience), 28 μl NaHCO_3_ (23 mg/ml), 28 μl 10x α-MEM and 4 μl of NaOH (1M) to adjust the pH to 10. The final collagen concentration of 1.2 mg/ml was prepared by diluting the initial solution with 310 μl of a mixture of 1 volume part of NaHCO_3_ (23mg/ml), 1 part of 10x α-MEM and 8 parts of distilled H_2_O. The pH was adjusted to 10 using NaOH (1 M). Cells were carefully mixed with the final collagen on ice to a final density of 4 × 10^4^ cells/ml and added to angiogenesis plates (Ibidi). The gel was formed by placing the sample in a cell culture incubator at 37°C, 5% CO_2_ and 95% humidity for 1 hour.

#### Encapsulation in Matrigel

A Corning® Matrigel® Matrix (ref. 356231) was used. 70 % Matrigel and 30% of cell solution with desired density were mixed on ice. The gel/cell mixture was then pipetted into the pre-coated Ibidi μ-slide, and incubated for 30 minutes at 37 °C to allow gelation.

### Fluorescence staining

At 24 h after encapsulation, gels were washed with PBS and then fixed with 4% paraformaldehyde (PFA) in PBS for 40 min. After fixation, the samples were permeabilized with 0.1% Triton X-100 in PBS for 10 min and blocked with 1% BSA in PBS for 30 min. The sample was incubated with Phalloidin Atto-520 (1:20 in 1% BSA/PBS, 10 μM, Sigma-Aldrich, USA) for 1 h for staining the actin fibers. After this, the cells were stained with Hoechst (40 μM) for 30 min and washed with PBS (3X). All procedures were performed at 37 °C. The images were acquired using Leica SPX8 microscope, equipped with a water objective (HC PL APO 63x/1.40, motCORR, Leica), and a HYD-SMD detector. As excitation sources, a supercontinuum White Light Laser (470–670 nm, pulsed, 80 MHz, NKT Photonics) and a UV diode laser (405 nm, pulsed, 40 MHz, Picoquant) were used. All measurements were performed in a temperature-controlled environment (37 °C).

### Rheology

#### Mechanical characterization of PIC gels

Both of the linear and nonlinear mechanical properties of PIC gels were characterized as previously reported^19^. Briefly, a stress-controlled rheometer (Discovery HR-2, TA Instruments) with an aluminum or steel parallel plate geometry was used (diameter = 40 mm, gap = 500 μm) to measure the mechanical properties of the hydrogels. Polymer solutions were loaded onto the rheometer plate at *T* = 5 °C, followed by a temperature ramp to *T* = 37 °C with a rate of 1.0 °C min^−1^. In the linear viscoelastic (LVE) regime, the moduli were measured at an amplitude of *γ* = 0.02 and a frequency of *f* = 1.0 Hz. Prior to the nonlinear measurements, the polymer solution was allowed to equilibrate for 30 minutes at 37 °C. Here, the hydrogels were subjected to a constant pre-stress of *σ*_0_ = 0.5 to 200 Pa, and the differential modulus *K′* was determined with a small superposed oscillatory stress at frequencies of *f* = 10 to 0.1 Hz (reported data at *f* = 1 Hz). The oscillatory stress was at least 10 times smaller than the applied pre-stress. In addition, creep and recovery tests were performed to characterize the plasticity of PIC gels. A constant stress ranging from 5 to 80 Pa was applied for 300– 3600 s while strain in response to the stress was measured over time. Then, the stress was removed in recovery tests and the strain was monitored until it reached equilibrium (Supplementary Fig. S10). The degree of plasticity is defined as the ratio of irreversible strain after recovery tests to the maximum strain at the end of the creep tests^45^.

#### Live cell rheology

A custom-made aluminum lower petri dish-like plate geometry (inner diameter 45 mm, height 10 mm) was designed and used to measure the mechanical properties of the cell-gel constructs. A cold cell-polymer solution (0.6 mL) was loaded into the dish center at 5 °C and the upper plate (20 mm) was lowered to a 1000 μm gap. Once the temperature was raised to 37 °C the gel containing the cells was allowed to stabilize for 5 minutes. Then warm CO_2_-independent medium (5 mL Gibco, Thermo Fisher, USA) was added into the dish. Next, a solvent trap cover was mounted to create a thermally stable vapor barrier. The storage modulus *G’* was determined by applying an oscillating deformation of amplitude *γ* = 2% with *f* = 1 Hz in a time sweep for 24-48 h. For the cell-gel rheology with CytoD experiments, 5 mL of 10 μM CytoD in warm α-MEM medium was added to the system during the pausing procedure at 24h, and then the measurement continued. PEG microspheres were kindly provided by Prof. Laura de Laporte (RWTH Aachen, Germany) with an average diameter of 20 μm. The seeding densities of live cells and PEG microspheres are both set to 10^6^/ml.

### 3D displacement microscopy

For cell-traction induced 3D matrix displacement measurements (3D displacement microscopy), adipose cells were incubated for 45 min with 12.5 μM CellTracker™ Green CMFDA (Life technologies, Belgium) in serum free α-MEM medium one day before gel encapsulation. PIC polymers were labeled with DBCO-Atto647N to fluorescently label the fiber network. The volume ratio between DBCO-Atto647N: cell solution: polymer solution was 1:9:10. Briefly, the dye (10 μM) and the polymer solution (2 mg/mL) were mixed on the ice first, and then the cold cell solution (80,000 cells/mL) was added and mixed gently. 12 μl of the mixture was pipetted in a μ-Slide Angiogenesis dish (Ibidi) and warmed up in an incubator to create cell-gel constructs. Collagen gel samples were imaged using second harmonic generation (SHG) microscopy^46,47^. For Matrigel samples, the gel deformation was track using embedded fluorescence beads (FluoSpheres™, 0.2 μm, red fluorescent (580/605), ThermoFisher, F8810). Similar cell densities were used for the different matrices.

#### Image acquisition

For PIC gel samples, fluorescence images were acquired using a Leica SP8 confocal microscope with a 25x water-immersion objective (NA 0.95) and a hybrid photomultiplier tube as detector (HYD-SMD, Leica). Cells labeled with CellTracker™ Green were excited at 492 nm, and PIC fibers grafted with Atto647N were excited at 646 nm with a non-sequential and bidirectional mode. The image stack was recorded with a distance of the confocal z-sections of 0.57 μm. With typically 175 z-slices and 512 x 512 pixels in the X-Y plane, the voxel dimensions were of 0.57 μm on the x, y and z direction. A series of time-lapse image stacks were recorded after adding CytoD (5 μM), with an interval time of 20 min between each z-stack. For collagen gel samples, fibers were imaged using 2-photon excitation at 808 nm with detection between 450 to 650 (550/200 nm BrightLine® single-band bandpass filter, Semrock). The other image acquisition settings were the same as for the PIC samples. A stage incubator was used to keep the cells were kept at 37 °C and 5% CO_2_ during image acquisition.

#### Displacement calculation

The Matlab toolbox TFMLAB was used to process the microscopy data and calculate matrix displacements around the adipose cell^29^. This software is freely available at: https://gitlab.kuleuven.be/MAtrix/Jorge/tfmlab_public. Briefly, the workflow of TFMLAB consists of three main steps: image processing, cell segmentation and displacement measurement.

##### Image processing

First, raw image data was filtered by penalized least squared-based denoising and enhanced with a contrast stretching operation. Second, stage drifts were corrected for by applying rigid image registration with respect to the relaxed state. The shift was calculated by means of a phase correlation operation on the fibril images (i.e. acquired by means of fluorescence imaging for the PIC hydrogels and by means of second harmonic generation for the collagen gels). The calculated shift was then corrected for on both the fibril images and the cell images.

##### Cell segmentation

Cell bodies were segmented by applying Otsu thresholding and by removing small binary objects (Supplementary Fig. S11a,b).

##### Displacement measurement and 3D visualization

TFMLAB uses the FFD-based image registration algorithm to register the stressed fibril images to the relaxed images. We used the normalized correlation coefficient as the similarity metric and a stochastic gradient descent method with adaptive estimation of the step size as the optimizer^47^. As a result, a 3D displacement field vector was obtained at every voxel of the image. The software Paraview was used to create 3D renders of the cells and the displacement field using the .vtk files provided by TFMLAB.

#### Displacement field analysis

To obtain the displacement decay away from the cell surface, we averaged the values of the displacement field magnitude at increasing distance from the cell surface with a step of 5 μm until the entire field of view was covered. Thus, an average displacement field value was obtained for each spatial bin, [0-5] μm, [5-10] μm…, and plotted in graphs shown (Fig. 2f, 4f and 5i).

### Cell morphology analysis

#### Automated 3D cell protrusion segmentation and cell morphology quantification

Morphology metrics were extracted from the cell binary mask provided by TFMLAB (Supplementary Fig. S11b). To automatically segment cell protrusions in 3D, an in-house algorithm was developed. First, the largest sphere that fits within the cell binary mask was calculated using Matlab functions *bwdist* and *pdist2* (Supplementary Fig. S11c). Letting R be the radius of the sphere, we selected the voxels from the cell binary mask located at an Euclidean distance larger than 2R. Connected components of this selection were taken as individual protrusions (Supplementary Fig. S11d). Protrusion length was calculated by measuring the principal axis length using Matlab function *regionprops3.* This function was also used to compute the volume and the solidity of the cells. In this work, we refer to the solidity of the cells as “sphericity” and expressed it as a percentage.

#### Automated cell segmentation of clumped cells

Multiple cells were present in low magnification images. In some cases, cells generated long protrusions and were in contact with neighboring cells. First, cell and nuclei images (Supplementary Fig. S12b,c) were filtered, enhanced and binarized as described above (Supplementary Fig. S12d). To accurately split the clumped cells, the watershed algorithm from the image processing toolbox dip image was applied using the nuclei masks as seeds. This provided the borders between cells in contact. After splitting the cells based on these borders, we obtained an individual mask for each cell (Supplementary Fig. S12e). Finally, we quantified the number of protrusions, sphericity and volume individually as described above.

### Matrix remodeling analysis

#### Image acquisition and processing

The images were acquired using the same settings as for TFM experiment. Cells were labeled with CellTracker™ Green, and PIC fibers were grafted with Atto647N. To quantify the degree of cell-induced fiber remodeling over time, the images were acquired at different time points (0, 2, 7, 15 and 24h) after encapsulation (Supplementary Fig. S13). To compare the matrix remodeling between stress and relaxes states, a series of time-lapse image stacks were recorded after adding cytoD (5 μM) with interval time 20 min (Supplementary Fig. S14). Furthermore, the raw image data was filtered by 3D Gaussian filtering.

#### Cell segmentation

The cell was segmented by applying Otsu thresholding and by removing small binary objects.

#### Polymer segmentation

The polymer was segmented by applying an intensity threshold based on the 99^th^ percentile intensity of the pixels on 10% outside frame of the image. Small objects were then removed, and small holes filled.

#### Calculations

The volume was calculated by summing all pixels inside the polymer segmented mask multiplied by the voxel size and subtracting the cell volume from it. The distance between the cell and polymer was calculated as follow: for each point on the cell mask surface, the closest point on the polymer mask surface (outer shell) was determined using Euclidian distances. We then looked at the distribution of those distances to have an estimate how far the densified region can extend from the cell.

#### 3D Modeling

The 3D models of cell and polymer were made by plotting the isosurface for i=0.5 which created a surface at the surface of the segmented mask.

#### Code source

All the code related to the matrix remodeling analysis can be found at: https://github.com/BorisLouis/fiberRemodeling

## Supporting information

**Fig. S1.**
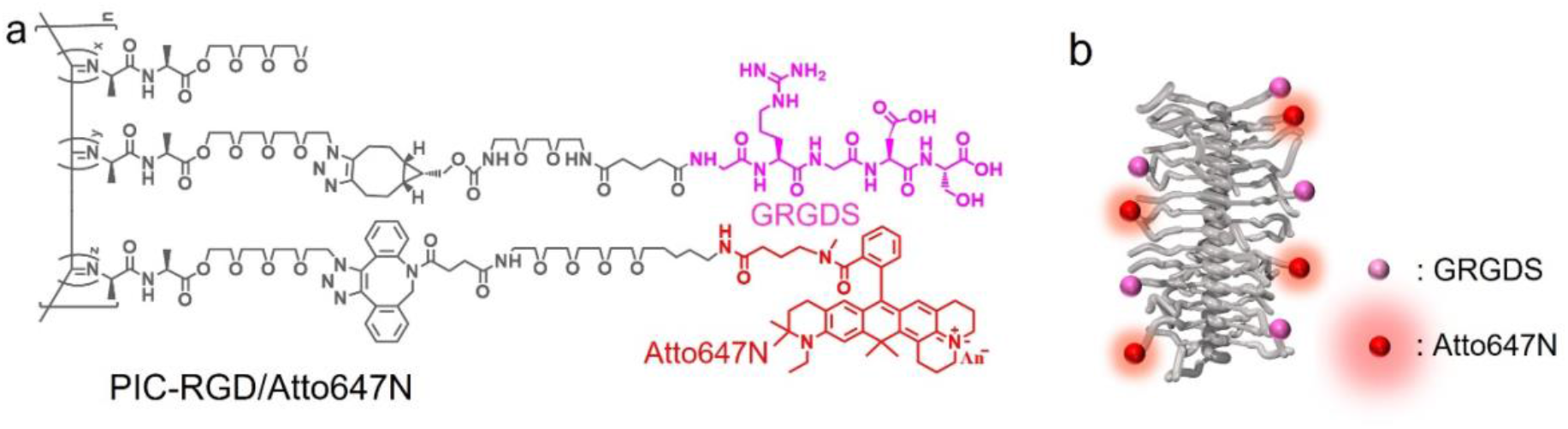
(a) Chemical structure of PIC polymers decorated with GRGDS and/or Atto 647N. (b) Schematic illustration of the helical structure of PIC molecules.

**Fig. S2.**
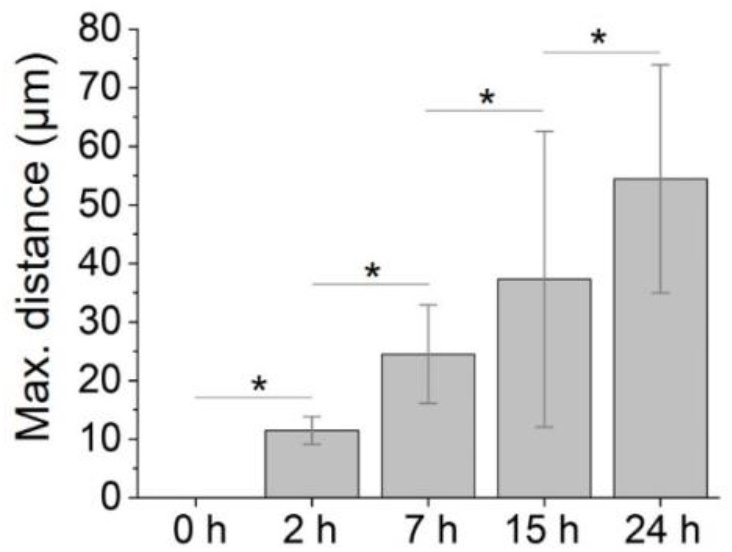
Maximum distance of the detected edge of fiber remodeling to cell surface at different time points of culture (N = 5, * indicates p<0.05, by One-way ANOVA Tukey’s test).

**Fig S3.**
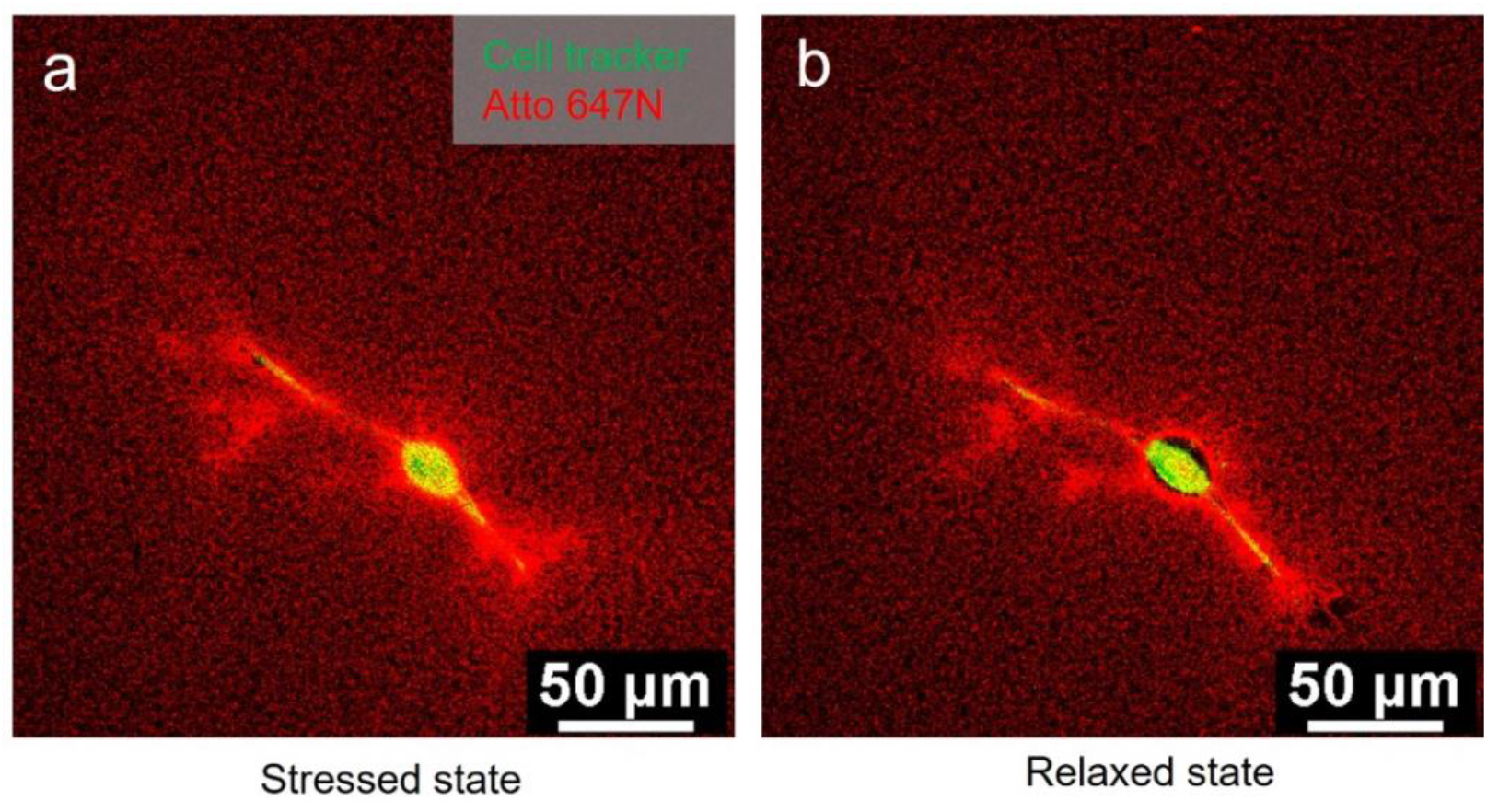
Schematic illustration of bead-free displacement microscopy approach based on PIC matrices. Representative fluorescence images of a cell (cell tracker in green) and surrounding PIC fibers (red) at the stress state (a), and the relaxed state (1h after adding cytochalasin D) (b).

**Fig. S4.**
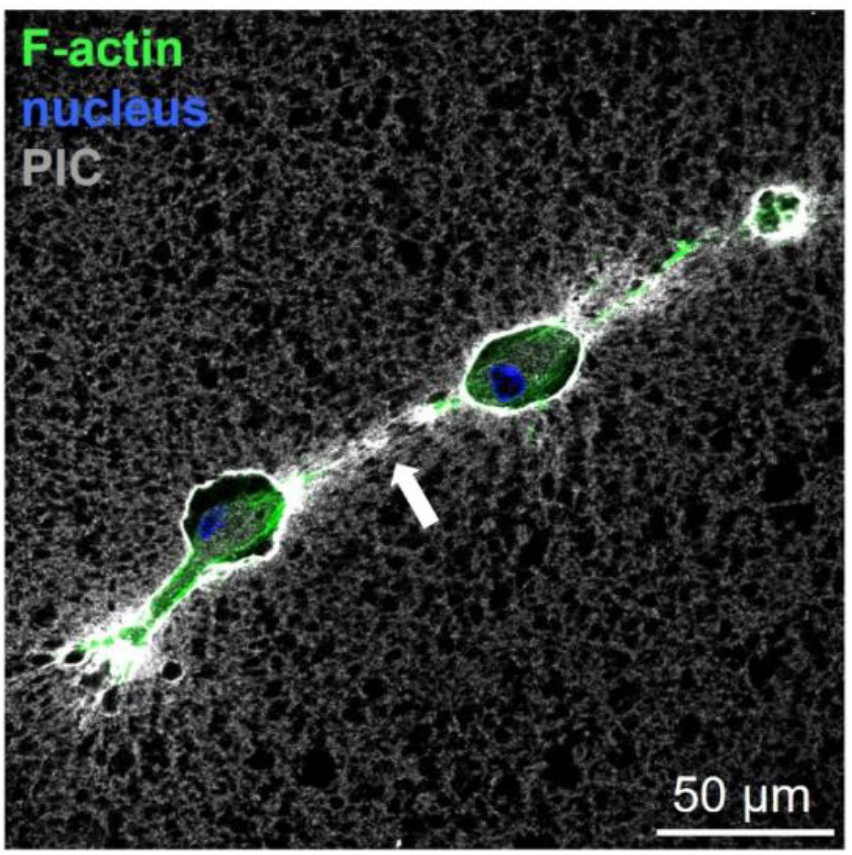
Two adjacent dipose cells in 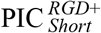 network, imaged 7 hours after encapsulation, showing fiber remodelling in the intercellular space. The remodelled fibres (white arrow) formed a track, enabling mechanical interaction between the cells. Scale bar = 50 μm.

**Fig. S5.**
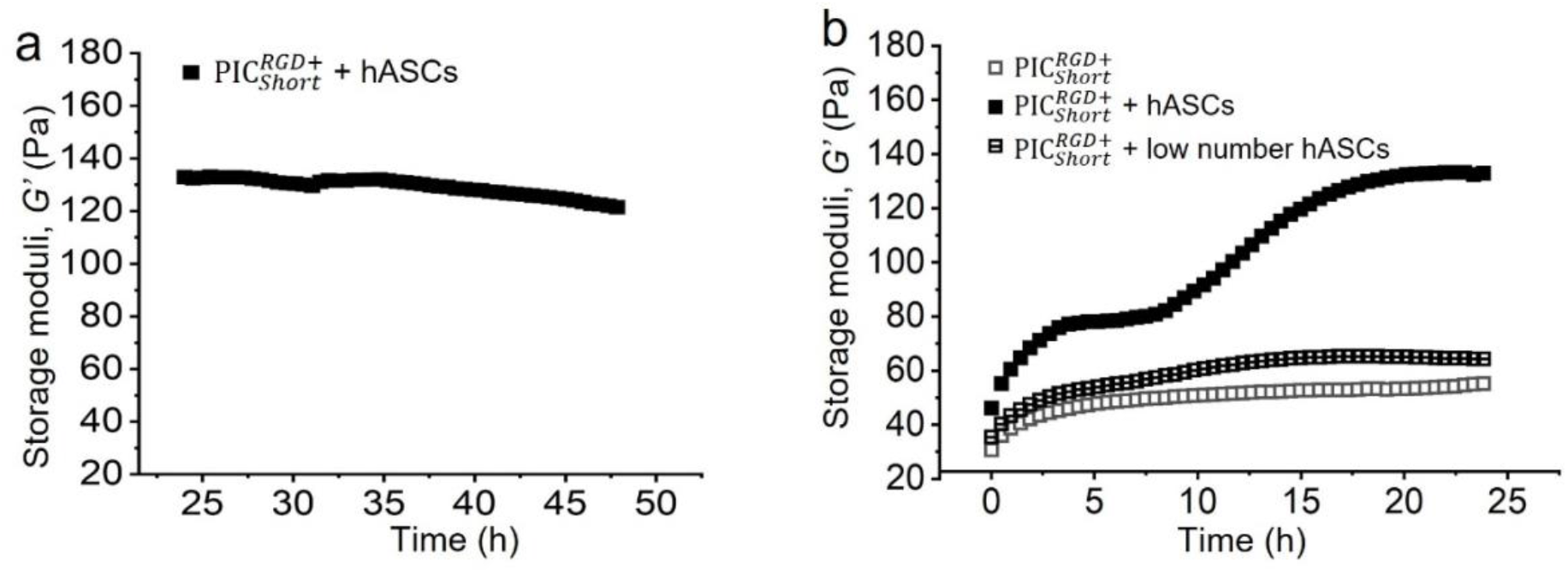
Impacts of variations of other parameters on gel stiffening. (a) The stiffness of the 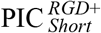 hASCs-gel construct remained constant during 24-48 hours after encapsulation. (b) Influence of the cell density on matrix stiffening in 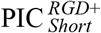. A low cell density (250,000 cells mL^−1^) was used. Note that in all gels, the polymer concentration was 1 mg mL^−1^, and the cell concentration was 10^6^ cells mL^−1^ unless otherwise specified.

**Fig. S6.**
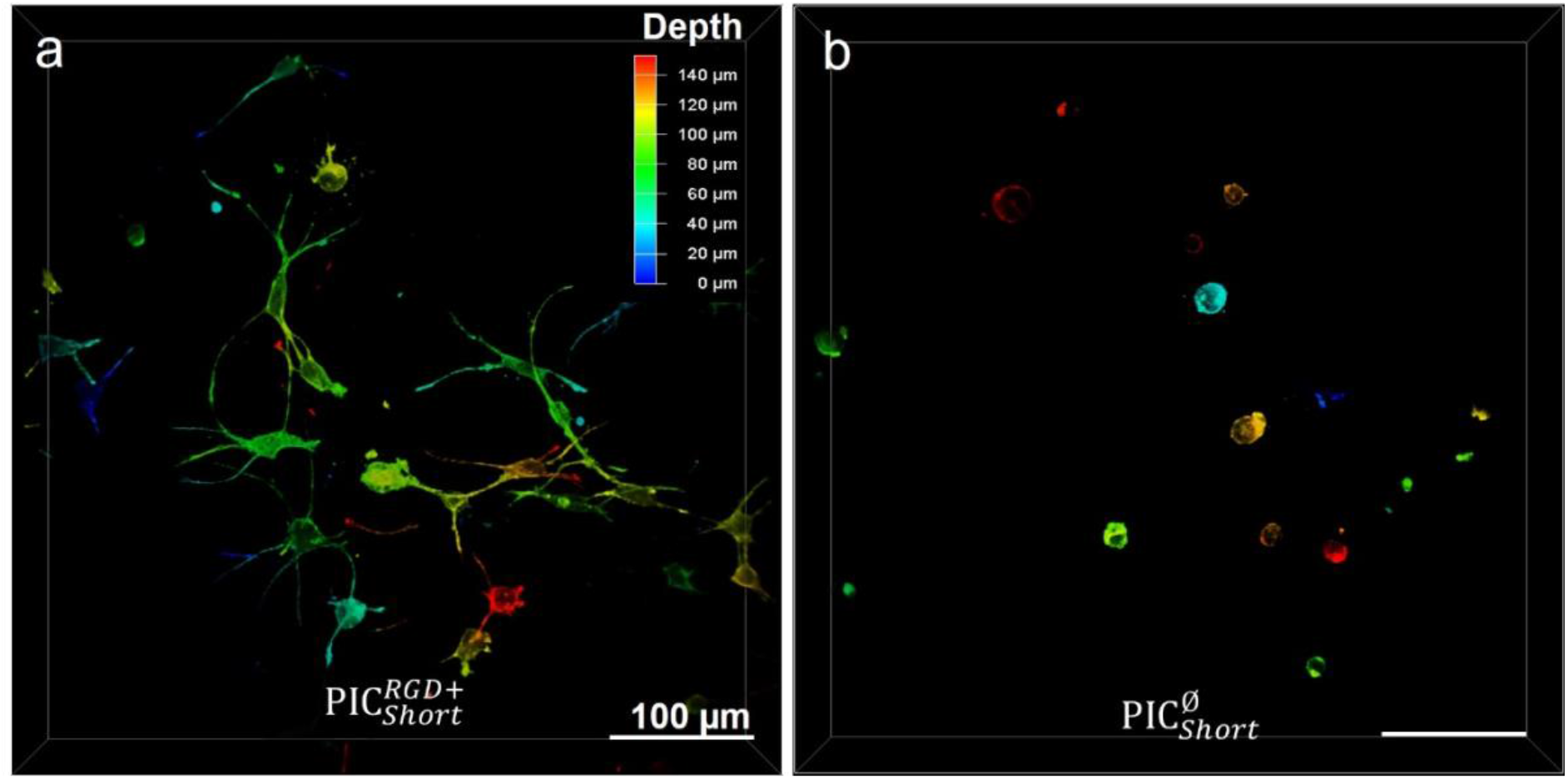
Color-coded (depth) 3D confocal fluorescence images showing cell morphology in PIC gels with (a) and without (b) RGD. Cells were labeled with phalloidin.

**Fig. S7.**
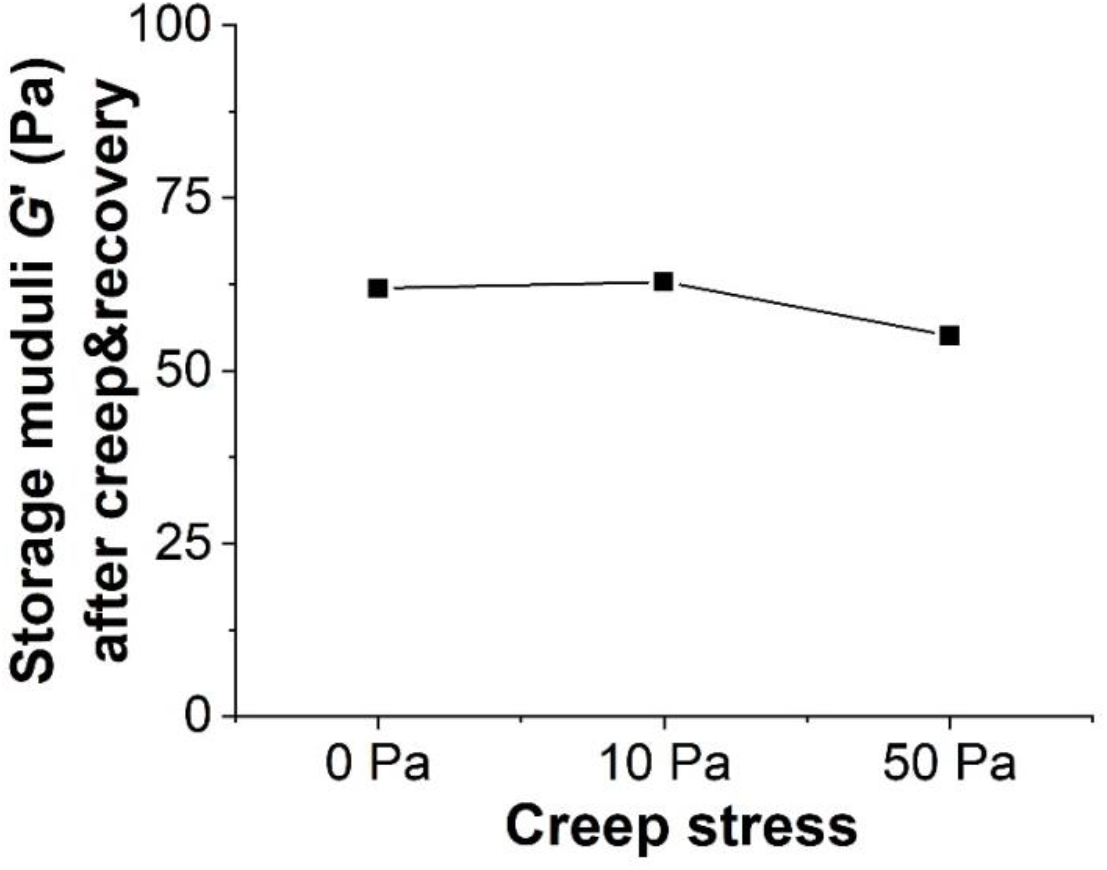
The shear storage moduli of the PIC gels 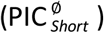 after a series of macroscopic creep and recovery tests with different applied stresses. The storage modulus G’ was determined by applying an oscillating deformation of amplitude *γ* = 2%, and *f* = 1 Hz. The polymer concentration was 1 mg mL^−1^.

**Fig. S8.**
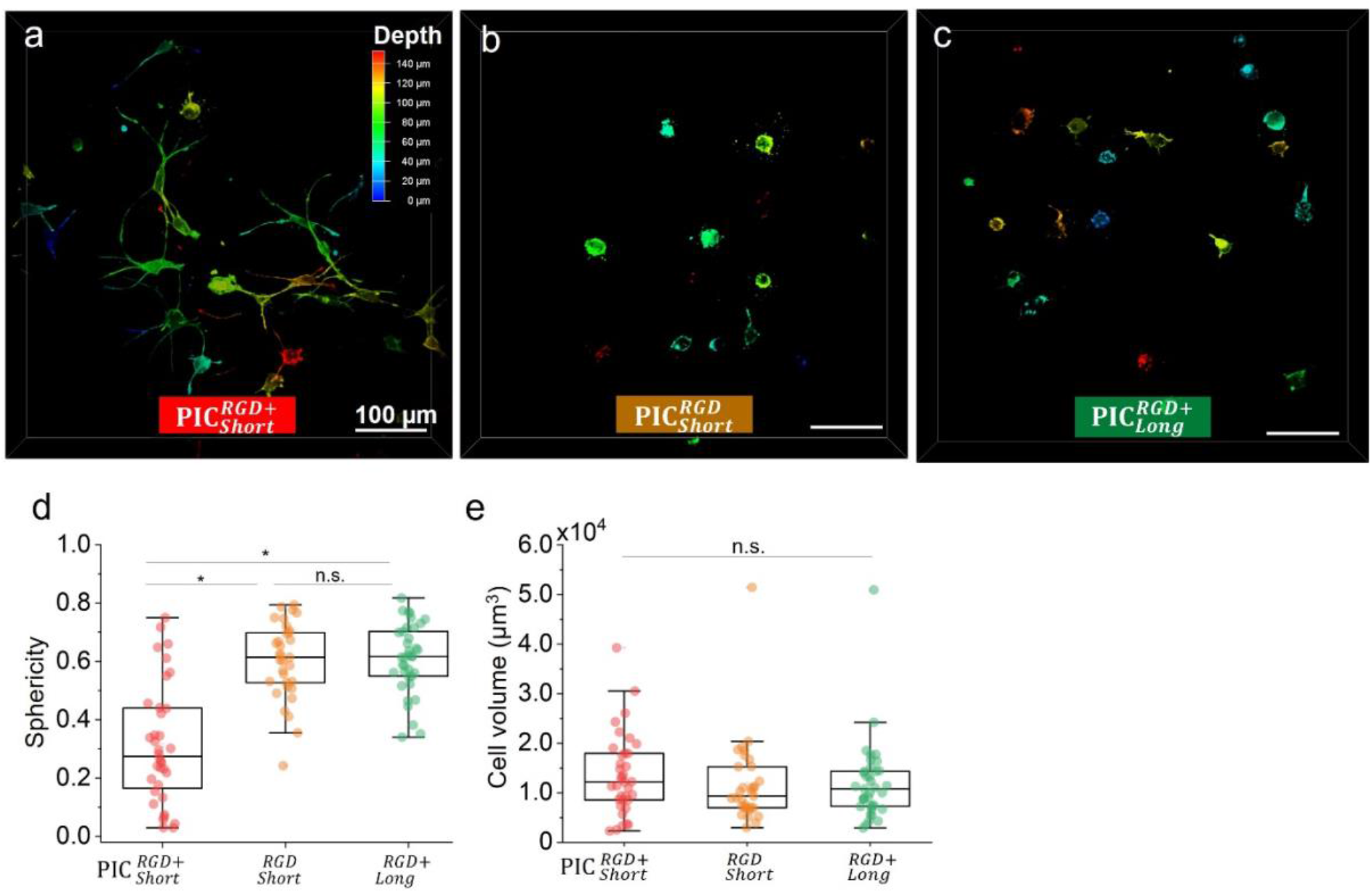
Color-coded (depth) 3D confocal fluorescence images showing cell morphology in PIC gels with different conditions 24 h after encapsulation, (a) 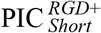, (b) 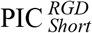, and (c) 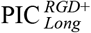. Cells were labeled with phalloidin. Analysis of cellular sphericity (d), and volume (e) in these three gels. Although cells show lower sphericity and higher number of protrusions in 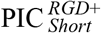 gels, protrusions are very thin and only take up a low percentage of the whole cell’s volume. Hence, a comparison based on cell volume does not reveal statistically significant differences

**Fig. S9.**
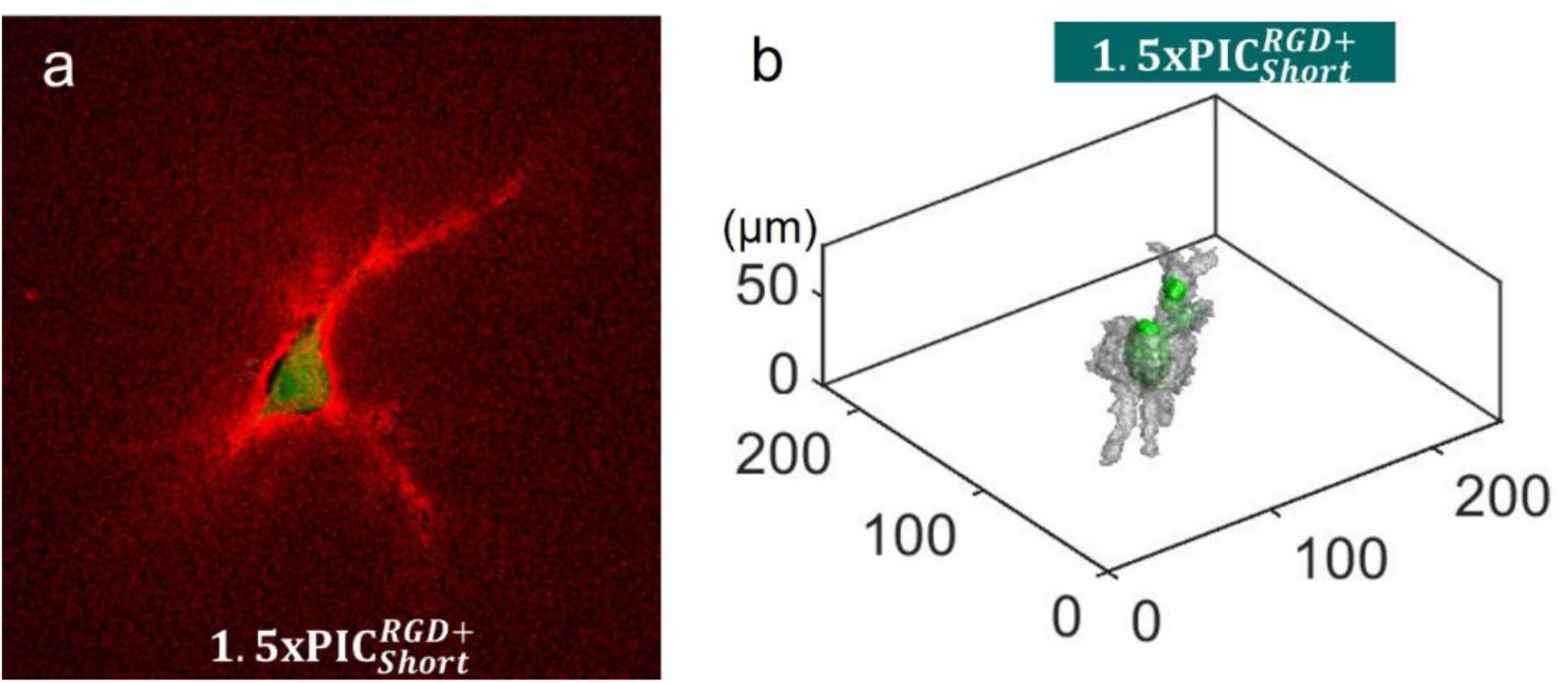
Investigation of the effect of linear stiffness on matrix remodeling. (a) Fluorescence images of cell induced remodeling 24 h after encapsulation in 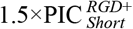 matrices (with a higher concentration at 1.5 mg/mL). Red: PIC fibers. Green: cells. (b) Representative 3D rendering of the remodelled matrix (grey) and the cell body (green).

**Fig. S10.**
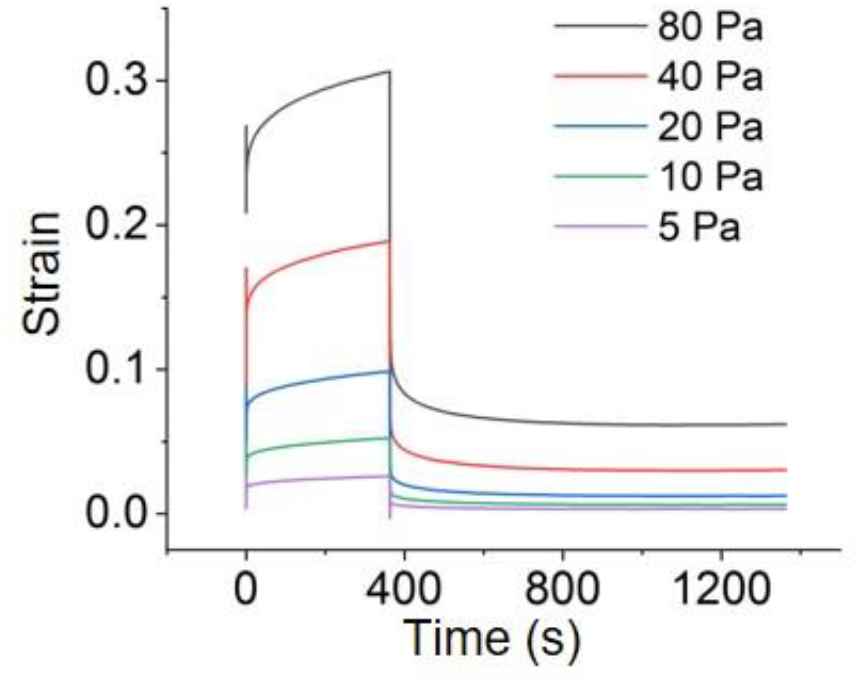
Creep and recovery tests characterize plasticity of PIC gels 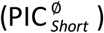. Initially, PIC gels exhibit instantaneous elastic response to applied stress, and strain of materials gradually increases over time. At the end of the creep test, PIC gels exhibit a maximum strain. After release of the creep test, the samples undergo elastic and viscoelastic recovery, but leave an irreversible strain, which is indicative of plastic deformation.

**Fig. S11.**
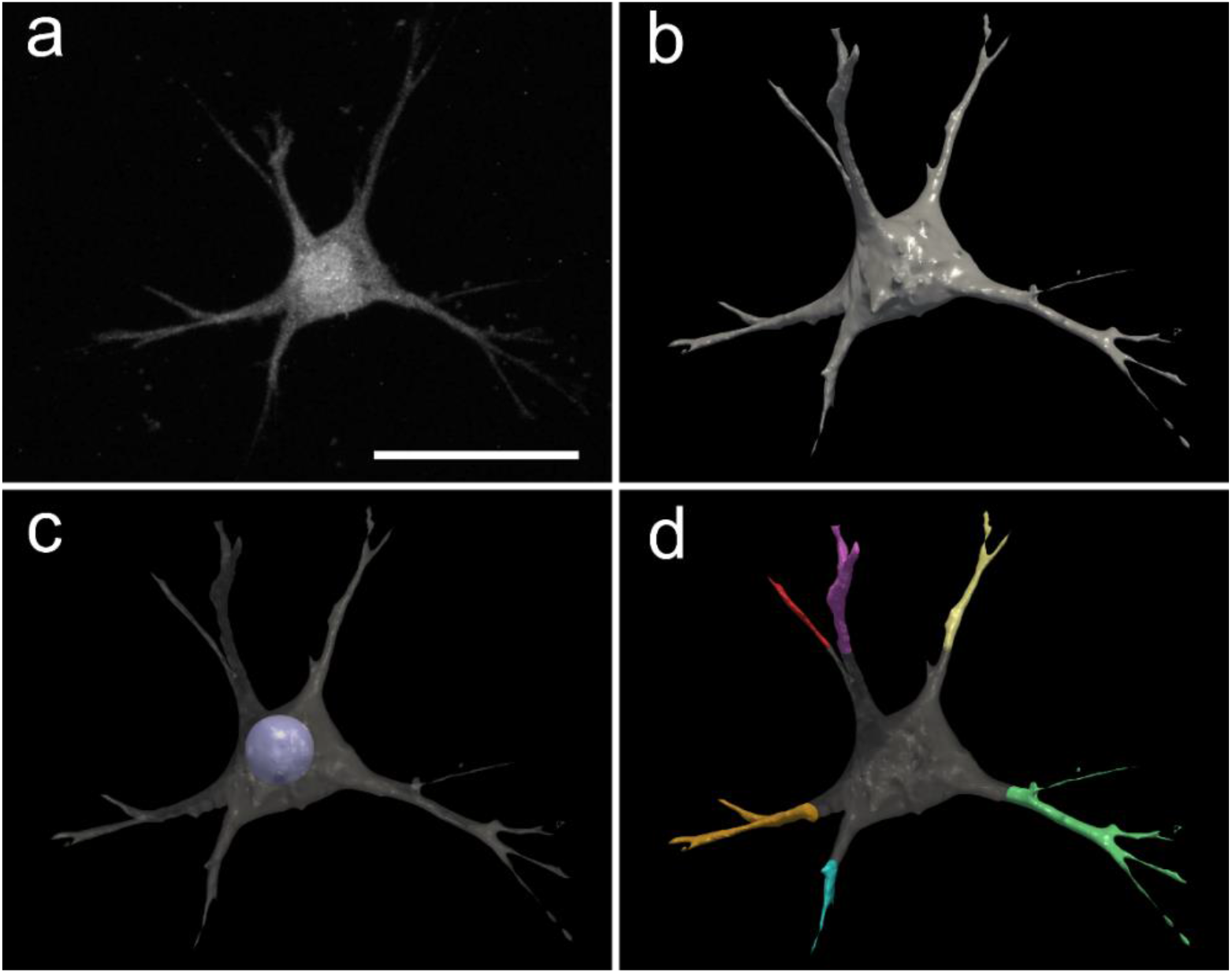
Illustrative example of the automated 3D cell protrusion segmentation. (a) Maximum intensity projection of an adipose cell in a PIC hydrogel acquired by means of confocal microscopy. Scale bar = 4.5 μm. (b) 3D binary cell mask. (c) Maximum sphere (purple) within the cell mask. (d) Segmentation of the cell protrusions. Each individual protrusion is labelled in a different color. b,c and d were rendered in Paraview.

**Fig. S12.**
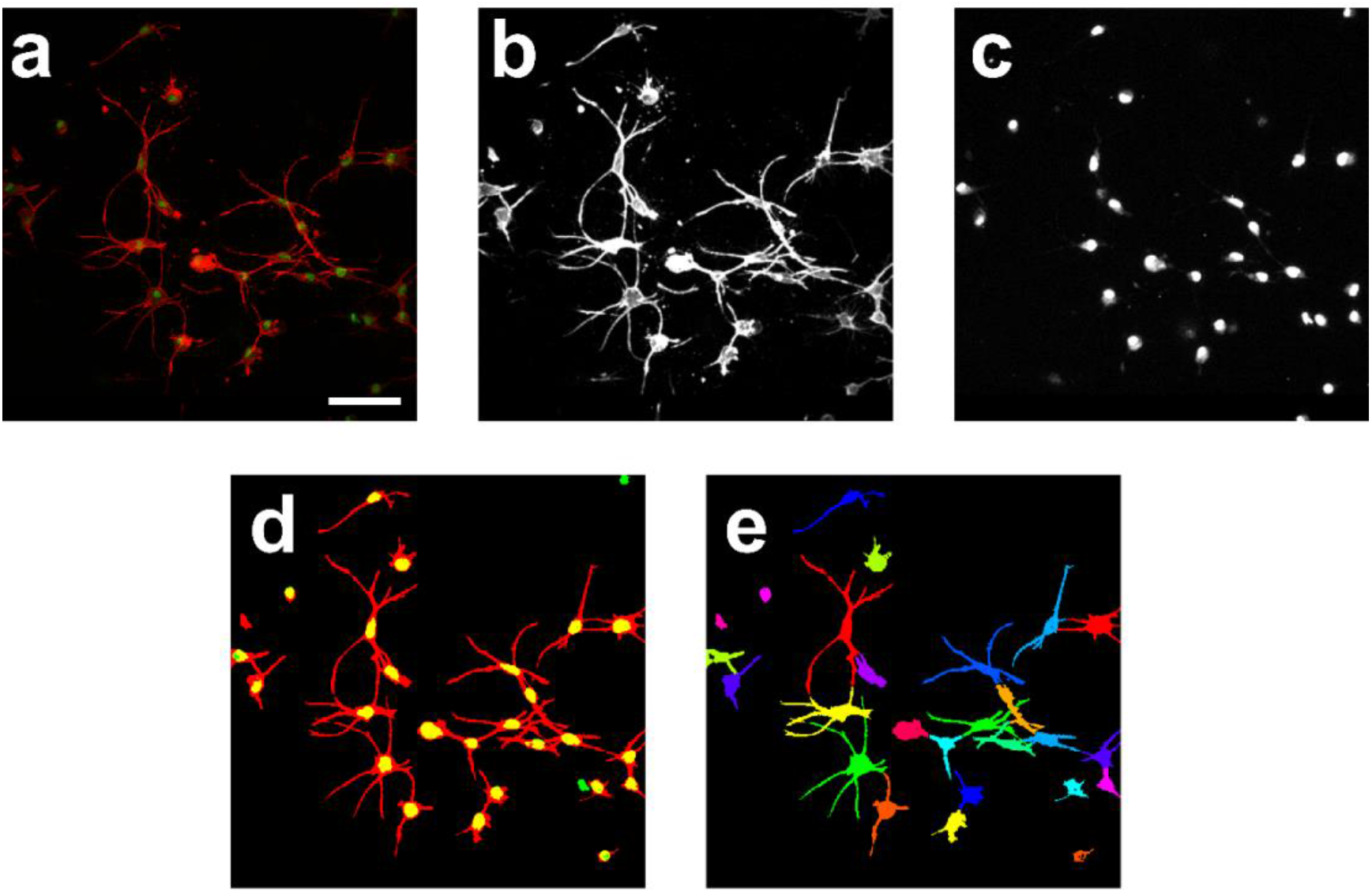
Illustrative example of the automated 3D cell segmentation of clumped cells. While the images shown in this figure are maximum intensity projections, the algorithm segments 3D cell volumes. (a) Raw data contain a cell channel (red) and a nuclei channel (green). (b) Filtered and enhanced cell channel. (c) Filtered and enhanced nuclei channel. (d) Binary masks of the cells (red) and the nuclei (green). Overlap is displayed in yellow. (e) Cell splitting after applying the watershed algorithm. Individual cells are labelled in different colors. Scale bar: 100 μm.

**Fig. S13.**
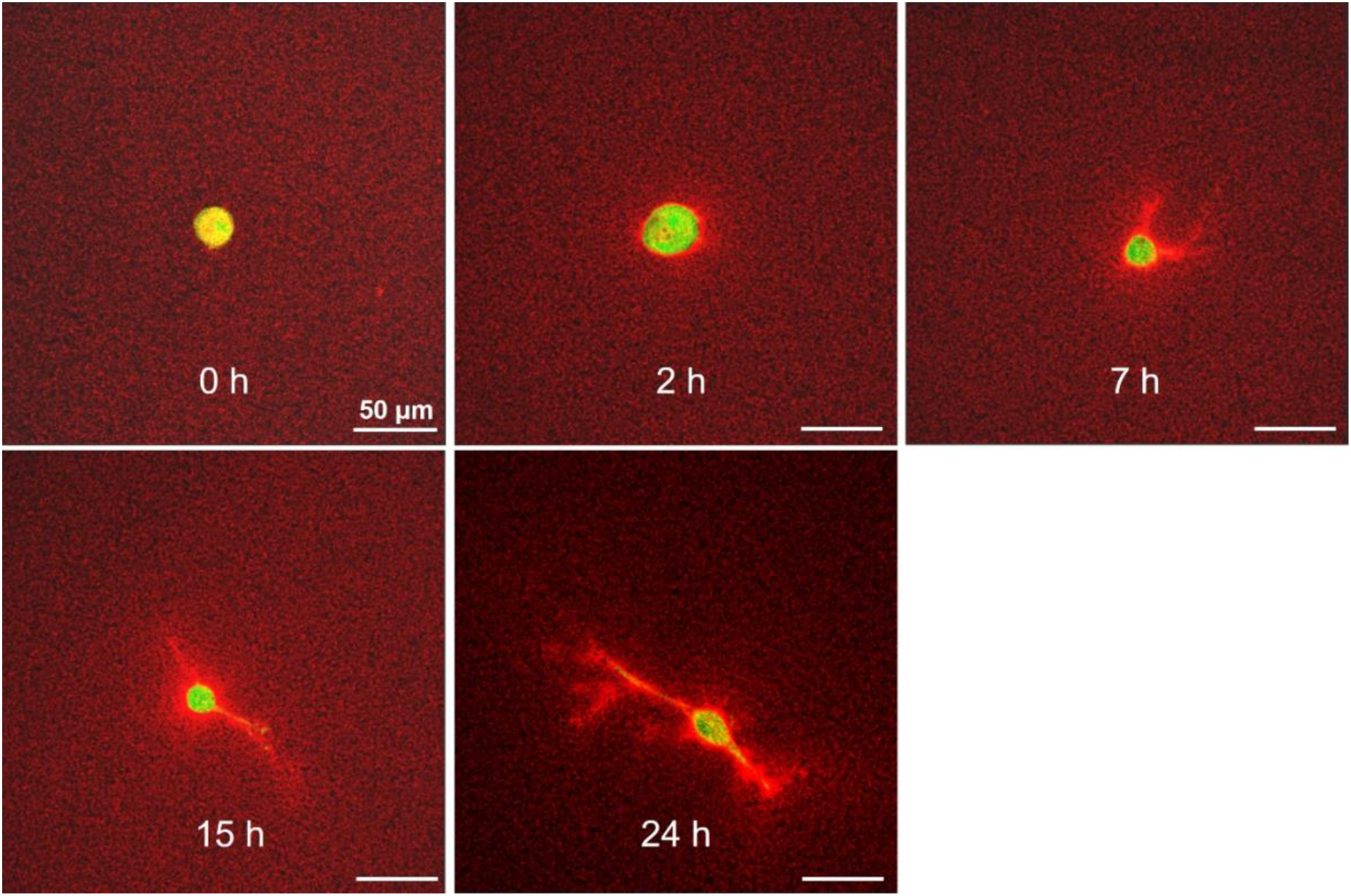
Fluorescence images of 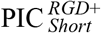 (1.0 mg/mL) with a representative encapsulated cell at different time points from 0 to 24 hours. Red: PIC fibers. Green: cells.

**Fig. S14.**
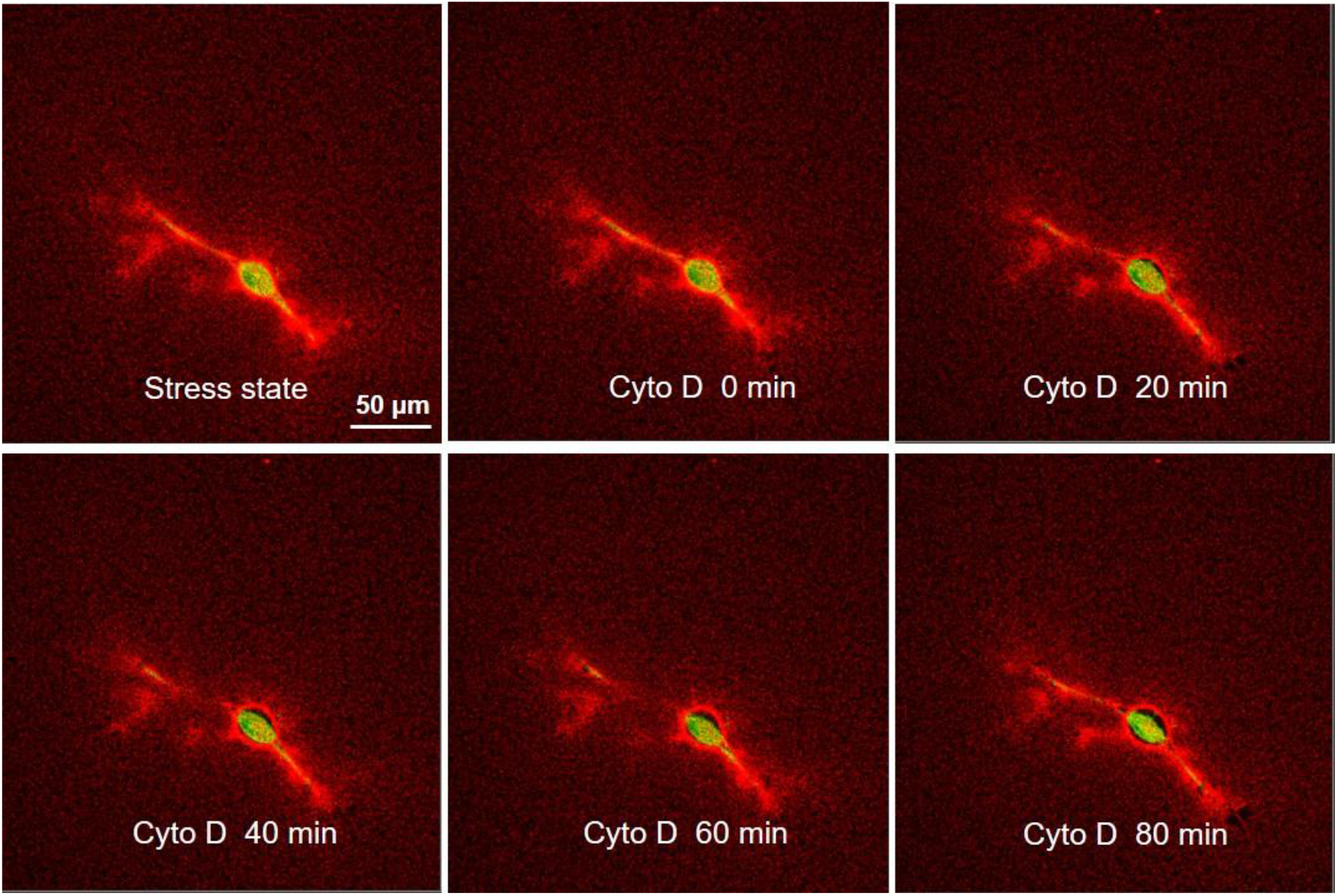
Representative time-lapse images show the changes of a fiber network (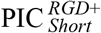, 1.0 mg/mL) and an encapsulated cell after adding Cyto D (5 μM). Red: PIC fibers. Green: cells.

